# A role for tubulin in cellular quality control and proteostasis

**DOI:** 10.64898/2026.04.06.716648

**Authors:** Sreya Basu, Nuo Yu, Riccardo Viscusi, Wouter Dof, Mirjam van den Hout, Wilfred F.J. van IJcken, Karel Bezstarosti, Dick H. W. Dekkers, Jeroen Demmers, Niels Galjart

**Affiliations:** Department of Cell Biology, Erasmus University Medical Center Rotterdam, The Netherlands; Department of Clinical Pharmacy, Erasmus University Medical Center Rotterdam, The Netherlands; Center for Proteomics, Erasmus University Medical Center Rotterdam, The Netherlands; Center for Biomics, Erasmus University Medical Center Rotterdam, The Netherlands

## Abstract

Microtubules, stiff rods built up from tubulin dimers, form a cytoskeletal network whose structure, behaviour, and function have been extensively investigated, mainly from a mechanical perspective. Here, we describe a role for tubulin in the cellular stress response. We overexpressed tubulin dimers in a controlled fashion in 293F cells. Despite the engagement of autoregulation, a mechanism that degrades tubulin-encoding mRNAs when tubulin levels are high, a surplus of tubulin and microtubules is detected in overexpressing cells. This leads to altered microtubule behaviour, mitotic problems, deregulation of the cell cycle, and replication stress. Surprisingly, we also observe proteostasis defects in tubulin overexpressing cells, which we attribute to mitochondrial stress-related translation attenuation. Conversely, tubulin and microtubules are downregulated as part of the response to oxygen or glutamine deprivation. Together, our data link tubulin levels, and hence autoregulation, to cellular quality control and proteostasis. We propose that competitive interactions with key partners, including the mitochondrial protein import and general translation machinery, underlie the tubulin-mediated control of cellular homeostasis.

---

Microtubules (MTs), one of the three major cytoskeletal networks of eukaryotic cells, are dynamic structures composed of heterodimers of *α* and *β* tubulin ^1^. These building blocks assemble in a head-to-tail arrangement into MTs, with one end of the MT (the minus end) exposing alpha-tubulin, and the other end (the plus-end) beta-tubulin. Soluble tubulin dimers are GTP-bound; they undergo GTP hydrolysis upon incorporation into MTs, resulting in a MT lattice that mainly consists of GDP-tubulin, and MT ends where GTP-tubulin prevails. In cells MT minus-ends are often embedded, whereas MT plus ends display stochastic cycles of polymerisation and depolymerisation, a behaviour called dynamic instability ^2^. The concentration of soluble tubulin dictates in part the rate of MT assembly, and thereby also the frequencies of catastrophe (the conversion of MT growth to shrinkage) and rescue (the conversion of shrinkage to growth), two major parameters of dynamic MT behaviour. MT growth, behaviour, and organisation are furthermore controlled by a large network of MT-associated proteins (MAPs) ^3^.

The *α* and *β* tubulin genes in mammals have been duplicated several times, giving rise to multiple highly homologous tubulin isotypes that are expressed in a tissue- and cell type-specific manner and that confer subtle properties to the MT they are part of ^4^. To simultaneously control isotype mRNA levels cells have a post-transcriptional mechanism in place, called tubulin autoregulation, in which imbalances in tubulin concentration at the protein level influence all the different tubulin-encoding mRNAs ^5^. Autoregulation was discovered in studies with MT-active agents (MTAs), compounds that bind tubulin on various sites ^6^ and that deregulate MTs either by stabilising the network or by de-stabilising it ^7^. The best-known MT stabiliser is paclitaxel (PTX), or taxol. In contrast, colchicine and nocodazole depolymerise MTs. It was found that colchicine treatment, which leads to a sudden rise in soluble tubulin levels in cells, causes the rapid degradation of tubulin mRNAs, whereas treatment with PTX leads to tubulin mRNA stabilisation ^5, 8, 9^. A key factor in autoregulation is the protein TTC5 which recognises nascent translating *α* and *β* tubulin polypeptides at the ribosome exit tunnel, and initiates mRNA degradation when tubulin levels are high ^10^. Recognition of the tubulin nascent chain requires a conserved N-terminus found on all tubulins (MREC on *α* tubulin and MREI on *β* tubulin). TTC5 bound to tubulin and the ribosome subsequently recruits SCAPER, an elongated protein that engages the CCR4-NOT deadenylase to degrade tubulin mRNAs ^11^. TTC5 is constitutively active in cells but its activity is low in normal conditions ^12^. Physiological functions for autoregulation remain poorly characterised.

While tubulin and MTs have been intensively examined from a mechanical perspective, roles for tubulin beyond MT assembly have been less well studied. We have previously established a system in which mammalian *α* and *β* tubulin are transiently co-expressed in equimolar amounts in cells, and we provided evidence that this “dual tubulin” approach gives rise to functional recombinant tubulins and MTs ^13^. Using the tagged dual tubulins, we performed affinity-based pull downs coupled to mass spectrometry-based proteomics and reported on a vast array of tubulin-associated proteins (TAPs) ^13, 14^. Here, we generated a Doxycycline (Dox)-inducible cell line to allow co-expression of *α* / *β* tubulin in a controlled manner. We examined MT behaviour and dynamics in cells overexpressing *α* / *β* tubulin, as well as the consequences of a mild tubulin overexpression. We find that overexpression generates surplus tubulin, despite the activation of tubulin autoregulation, as well as an excess of MTs. This perturbs MT behaviour and multiple processes in cells, including the cell cycle, which results in genome maintenance problems. Surprisingly, tubulin overexpression also induces mitochondrial dysfunction, skews the proteome by translation attenuation, and affects other stress responses. Conversely, amino acid and oxygen deprivation in normal cells suppress tubulin and the MT network. Our data reveal an important role for tubulin in cellular homeostasis.

## Results

### Characterisation of an inducible 293F cell line overexpressing α- and *β*-tubulin

We used our “dual tubulin” strategy ^13^ to produce novel human recombinant *α* and *β* tubulins from a single mRNA. Instead of the relatively bulky GFP moiety used previously, we placed a 6xHIS tag upstream of alpha-tubulin, and we added two TEV protease cleavage sites, one upstream of P2A and one downstream of the HIS-tag (Fig. 1a). Recombinant TUBB3 (rTUBB3) and TUBA1A (rTUBA1A), which can dimerise with each other and with endogenous tubulins (Fig. 1a), are distinguished on western blot (Fig. 1b), and their levels with respect to endogenous tubulins can therefore be easily quantified.

**Figure 1.**
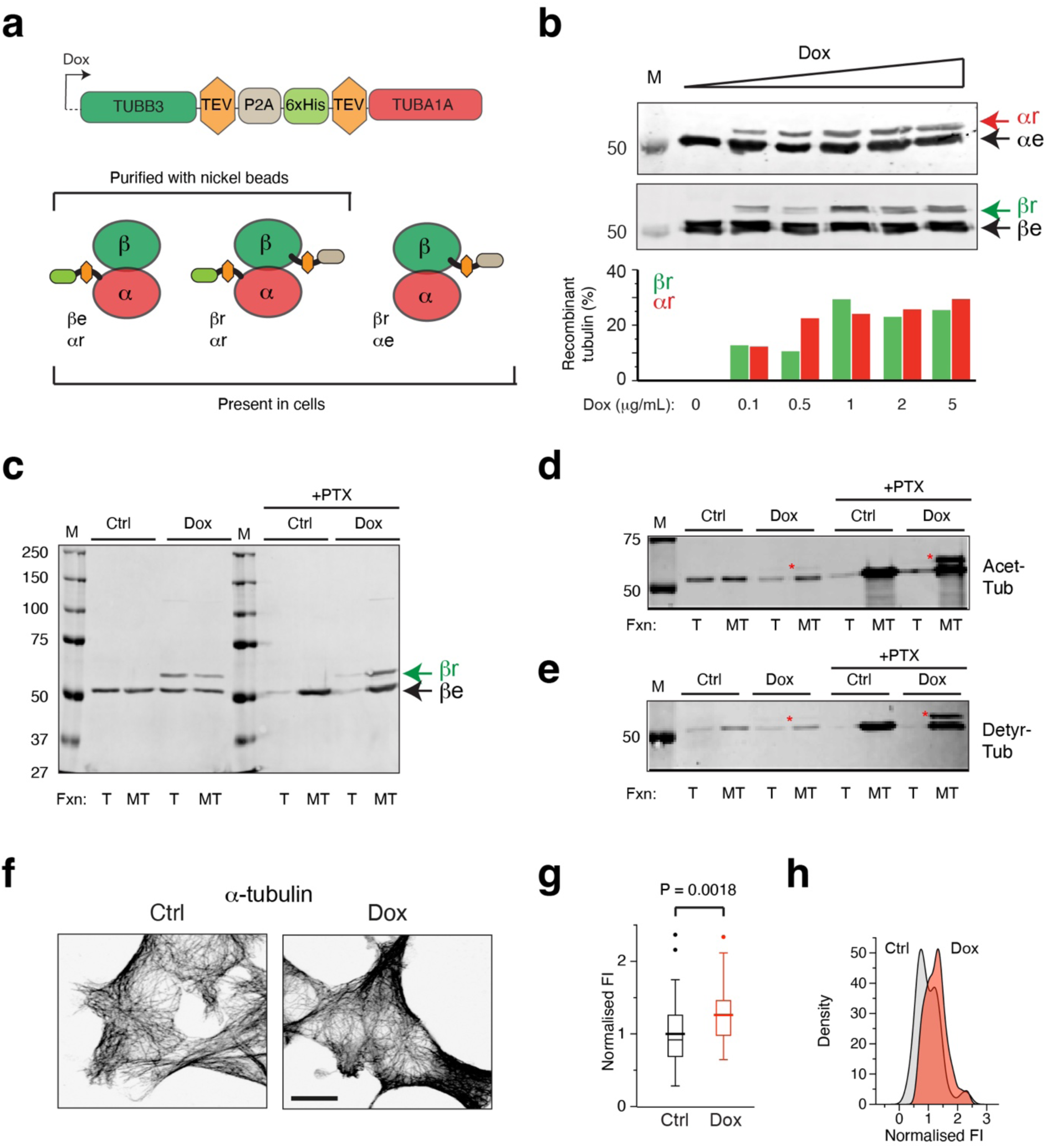
Inducible expression of recombinant *α*/*β* tubulin. **a)** Schematic representation of dual recombinant tubulin construct and tubulin dimer configurations. Upper panel: dual recombinant tubulin construct expressed under Tet-responsive promoter. TEV: TEV protease cleavage site, P2A: the P2A self-cleaving peptide sequence, 6xHis: six consecutive histidines. Lower panel: recombinant alpha- and beta-tubulin subunits dimerise with endogenous tubulins and with themselves generating three types of tubulin dimers in cells with recombinant tubulin. Dimers containing recombinant alpha-tubulin are purified with nickel beads. **b)** Controlled overexpression of recombinant tubulins. Representative western blots showing recombinant and endogenous tubulin levels in increasing Doxycycline (Dox) concentrations. Cells were harvested after 24 hours of Dox-induction. Upper panel: western blots (*β*r, recombinant beta-tubulin, *β*e, endogenous beta-tubulin, αr, recombinant alpha-tubulin, αe, endogenous alpha-tubulin, coloured arrows indicate the recombinant proteins). Lower panel: quantification of recombinant tubulin levels, the quantity of endogenous tubulin signal is set at 100%. **c-e**) MT fractionations in cells. Cells were either treated with Doxycycline (Dox) for 24 hr or not treated (control: Ctrl). Where indicated paclitaxel (PTX) was added at a concentration of 10 µM for 2 hours. Cell lysates were analysed by SDS-PAGE and western blotting. In (**c**) anti-beta-tubulin antibodies were used, in (**d**) and (**e**) antibodies against acetylated tubulin (**d**) or detyrosinated tubulin (**e**) were used. Red asterisks mark recombinant acetylated (**d**) and detyrosinated (**e**) tubulin. Representative blots shown from 3 independent experiments. **f**) MT distribution in 293F cells. Cells were treated with Dox for 24 hours, or not treated (Ctrl). Cells were fixed, and stained with anti-alpha-tubulin antibodies. Representative immunofluorescence stainings are shown. Scale bar: 10 µm. **g, h**) Quantification of intracellular MT density. Regions of interest (ROIs) were placed in IF images as shown in (**f**) and average fluorescence intensity (FI) was measured. Panel **g** shows a whisker plot of the normalised FI, mean values are indicated by the wider lines. Panel **h** shows a histogram of the density of the normalised FI. Control: n= 49 Dox: n=46, 3 independent experiments. Student T-Test reveals significant difference between Ctrl and Dox.

The dual construct was placed downstream of a Tet-responsive element, and transfected into human 293F cells engineered to express a transgenic reverse-Tet transactivator (rTTA) gene ^15^. We generated multiple clones containing a stably integrated construct and selected one line that expressed rTUBB3 and rTUBA1A up to ∼30% of endogenous tubulin levels (Fig. 1b). To examine functionality, we isolated recombinant tubulin using Ni-NTA affinity chromatography (Fig. S 1a, b). We obtained purified rTUBA1A, dimerised with rTUBB3 as well as endogenous *β*-tubulin (Fig. S 1c). We then performed MT polymerisation assays, at different temperatures and with or without PTX. In the presence of PTX more than 90% of all tubulin (recombinant and endogenous) was found in the MT pellet, whereas at 4^0^C hardly any tubulin polymerised (Fig. S 1d), demonstrating that purified recombinant tubulin is assembly-competent.

To investigate recombinant tubulin properties in cells we fractioned cell lysates using high speed centrifugation. Recombinant tubulin was detected both in the soluble (tubulin-containing) and pellet (MT-containing) fractions (Fig. 1c). Addition of PTX for two hours prior to cell isolation greatly increased the amount of MTs, which also contained recombinant tubulin (Fig. 1c). Furthermore, two common post-translational modifications (PTMs) on alpha-tubulin in MTs, i.e. acetylation at the K40 residue and detyrosination (removal of the terminal Y residue) ^16–19^, were observed on recombinant α-tubulin (Fig. 1d, e, respectively). We detected an increase of these PTMs on recombinant tubulin after pre-treatment of cells with PTX (Fig. 1d, e), which has been shown to induce MT acetylation and detyrosination ^20^. Taken together, these data suggest that recombinant tubulins are functional in cells.

We next examined the interphase MT network in Dox-induced and control 293F cells using immunofluorescence microscopy. Cells were grown on coverslips, fixed with methanol (which largely removes soluble tubulin), and stained with antibodies against α-tubulin (Fig. 1f). Fluorescence intensity of the secondary antibody was taken as a readout of MT density. We observed a 1.3-fold higher fluorescence intensity in Dox-induced cells compared to control cells (Fig. 1g), similar to the increase in soluble tubulin. The whole cell population contained excess MTs (Fig. 1h), indicating that Doxycycline induces an excess of tubulins and MTs in all cells. This situation is unique compared to cells treated with high concentrations of MTAs, which either lead to an increase in MT mass with concomitant loss of soluble tubulins (e.g. PTX treatment), or to an increase in soluble tubulins and a loss of MTs (e.g. nocodazole or colchicine treatments).

### Tubulin overexpression affects dynamic MT behaviour

To assess the effect of dual tubulin overexpression on MT dynamics we transfected control and Dox-induced cells with a cDNA encoding the MT plus-end marker EB3-GFP ^21^. We acquired time-lapse videos of EB3-GFP 24 hr after transfection (Fig. 2a, Supplementary Videos 1, 2) and examined MT growth rates as well as growth duration (the inverse of catastrophe frequency). We detected a mild increase in MT growth rates in Dox-treated cells (Fig. 2b), which was observed in all of the growing MTs examined (Fig. 2c), consistent with an increase in soluble tubulin levels in all Dox-induced cells. Surprisingly, a slight decrease in growth duration was detected (Fig. 2d), indicating that although MTs grow faster after Dox addition the rate of catastrophe is slightly higher.

**Figure 2.**
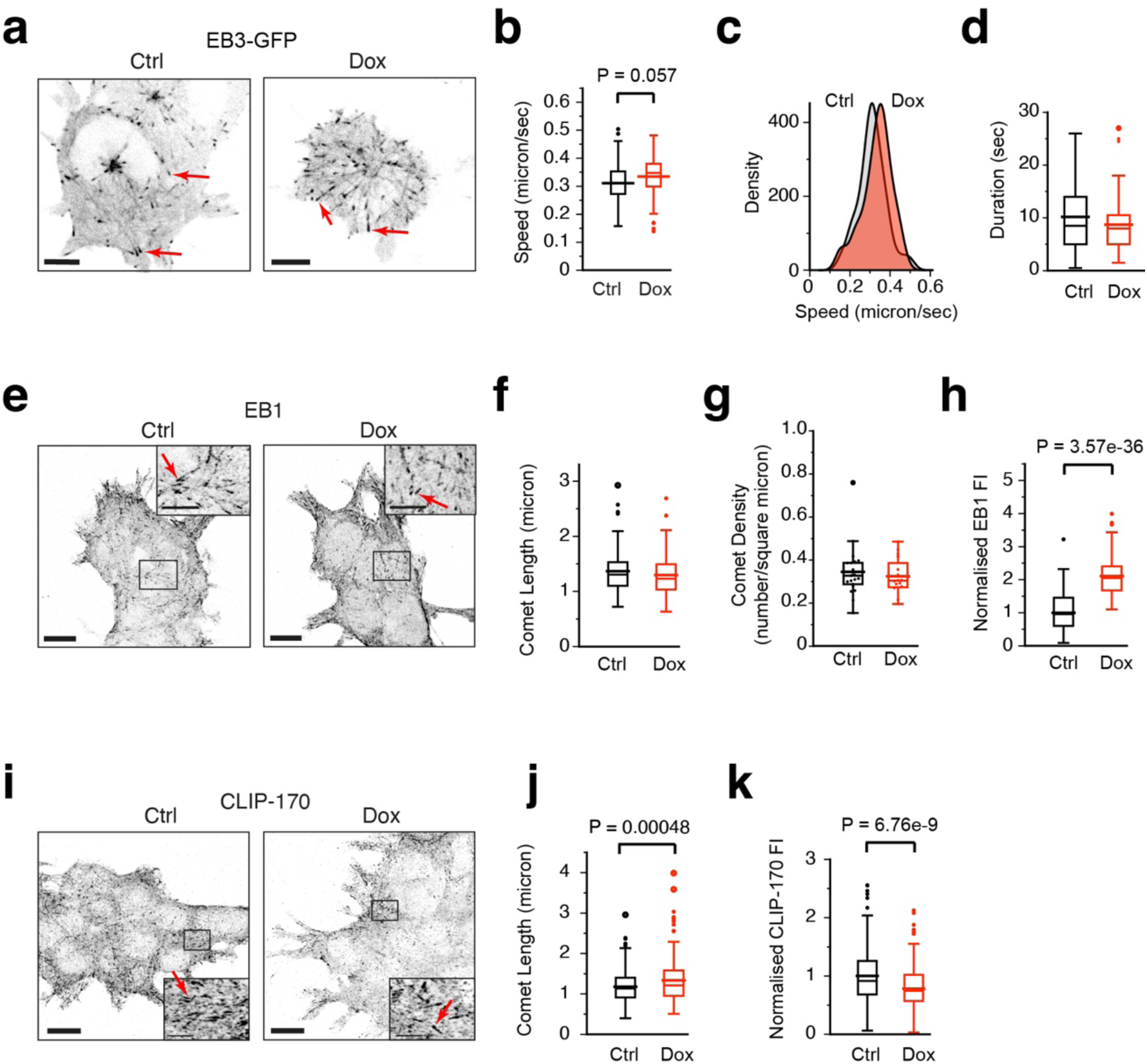
Tubulin overexpression affects dynamic MT behaviour. **a)** EB3-GFP localisation in transfected 293F cells. Cells were treated with Doxycycline (Dox) for 24 hours, or not treated (Ctrl). Examples of EB3-GFP comets are indicated with red arrows. Scale bar: 5 *μ*m. **b-d**) Quantification of dynamic EB3-GFP behaviour. EB3-GFP displacements in time, which represent movement of growing MT ends, were analysed in EB3-GFP-transfected 293F cells by spinning disk microscopy. 293F cells were treated with Dox for 24 hours, or not treated (Ctrl). We analysed 41 ctrl and 51 dox-treated plus-ends from 3 independent experiments. (B) shows a whisker plot of the speed, with mean values also indicated (wider lines). (C) shows a histogram of the density of the normalised FI. (D) shows a whisker plot of the duration, with mean values also indicated (wider lines). **e**) EB1 localisation in 293F cells. Cells were treated with Dox for 24 hours, or not treated (Ctrl). Cells were fixed and stained with anti-EB1 antibodies. Shown is a maximum intensity projection of 2 z-slices. Examples of EB1 comets are indicated with red arrows. Scale bar: 10 *μ*m, inset 5 *μ*m. **f-h**) Quantification of EB1 localisation. Fluorescence images as shown in (E) were analysed for EB1 comet length (**f**), comet density (**g**), and fluorescence intensity (FI) of comets (**h**). Shown are whisker plots, with mean values also indicated (wider lines). In (H) fluorescence images as shown in (E) were analysed for EB1 comet density by counting the number of plus-ends in defined ROIs for each image. For F,G, n= 132 control and 129 Dox-plus-ends, 3 independent experiments. For H, 21 ROIs for ctrl and 16 for Dox-treated cells, from 2 independent experiments Significant differences were examined using T-Tests. **i**) CLIP-170 localisation in 293F cells. Cells were treated with Dox for 24 hours, or not treated (Ctrl). Cells were fixed and stained with anti-CLIP-170 antibodies. Examples of CLIP-170 comets are indicated with red arrows. Scale bar: 10 *μ*m, inset 2.5 *μ*m. **j, k**) Quantification of CLIP-170 localisation. Fluorescence images as shown in (I) were analysed for CLIP-170 comet length (J) and fluorescence intensity (K). Shown are whisker plots with mean values also indicated (wider lines). n=202 ctrl and 155 Dox-treated plus-ends from 2 independent experiments. Student T-Tests reveal significant differences between Ctrl and Dox.

We next examined the localisation of the +TIPs EB1 and CLIP-170. Immunofluorescence (IF) staining for EB1 (Fig. 2e) did not reveal differences with respect to comet length (Fig. 2f) or overall number of plus-ends per unit area (Fig. 2g). However, the fluorescence intensity of EB1 at the MT end was approximately 2-fold higher in Dox-induced cells (Fig. 2h), indicating that twice as many EB1 proteins are bound per MT end in Dox-induced cells. By contrast, IF stainings for CLIP-170 (Fig. 2i) revealed an increased CLIP-170 comet length in Dox-induced cells (Fig. 2j), whereas fluorescence intensity at the MT end was decreased (Fig. 2k).

Combined, our results demonstrate that mild overexpression of tubulin alters MT growth rate and catastrophe frequency, and MT end behaviour. It has recently been shown that an increase in the amount of the TUBB3 isotype in cells is linked to a decrease in the lifetime of MT growth (i.e. increased catastrophe frequency) and a shorter region of EB binding at the plus end ^22^, indicative of an altered GTP cap ^23^. The higher level of recombinant TUBB3 in Dox-induced cells therefore explains the increased catastrophe frequency that we observed. However, we also detected twice as much EB1 per MT end, yet less CLIP-170 molecules, which were instead distributed over a larger region. We ^13^ and others ^24^ have shown that tubulins interact with CLIP-170. Conversely, CLIP-170 binds tubulin ^25–27^. An increase in the amount of soluble tubulin in Dox-induced cells may sequester CLIP-170 in the cytoplasm, causing a reduced accumulation of this +TIP at MT ends. Alternatively, an increase in soluble tubulin may affect liquid-liquid phase separation properties of +TIPs, and hence their plus-end distribution ^28–30^. While the molecular mechanisms need further investigation, our results conclusively show that soluble tubulin levels control +TIP composition at MT ends.

### Mitotic defects in 293F cells overexpressing tubulin

MTs are crucial players in mitosis, as they form the mitotic spindle, that allows chromosomes to be segregated ^31^. Defects in MT behaviour during mitosis can lead to chromosome misalignment, and activation of the spindle assembly checkpoint (SAC), which delays chromosome segregation until mitotic errors are resolved. Although the SAC is a potent mechanism, it can be overcome, and cells eventually divide despite segregation errors, leading to chromosome instability (CIN) and cancer ^32, 33^. Since we observed an increased tubulin and MT content in Dox-induced cells, accompanied by altered MT behaviour and +TIP composition at MT ends, we examined mitosis in these cells. We first followed MTs in live cells, using Sir-Tubulin to stain MTs ^34^. Light sheet fluorescence microscopy (LSFM), which allows for fast acquisition of large fields of view with low levels of photobleaching ^35^, revealed that the average time taken to complete metaphase was higher after Dox-induction (Fig. 3a, b) indicating that the SAC is activated in Dox-induced cells to delay anaphase onset, similar to results with MTAs ^36–38^. We also observed “extra-mitotic” centrioles at the onset of mitosis after Dox-treatment, indicative of mitotic arrest and aneuploidy, which eventually resolved into normal metaphase plates (Supplementary Videos 3, 4). When we examined mitotic structures in fixed cells, we observed increased chromosome mis-alignment at the metaphase plate (Fig. 3c, d), indicating that chromosome segregation was impaired in Dox-induced cells. Combined, these data point to activation of the SAC upon tubulin overexpression.

**Figure 3.**
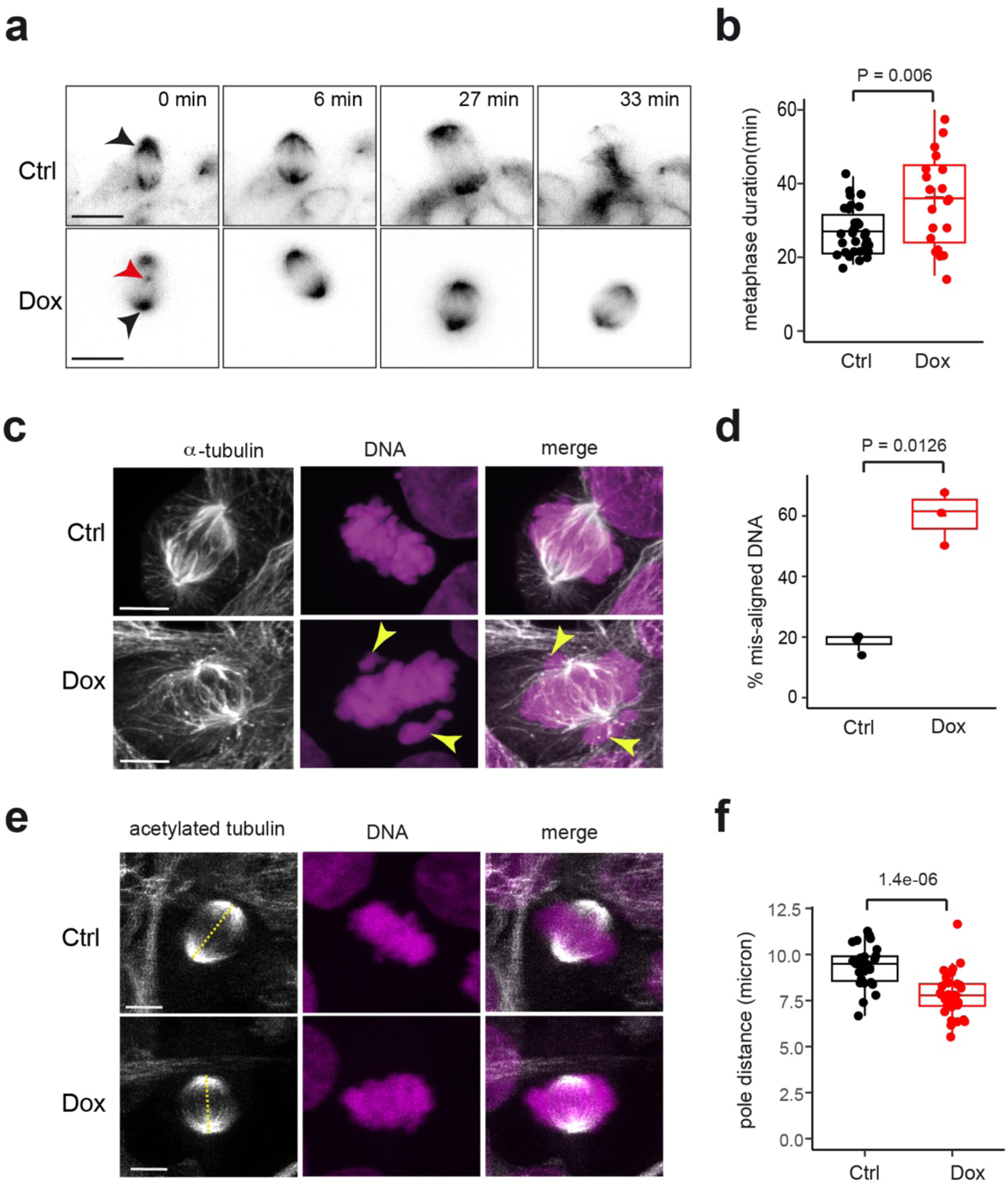
Tubulin overexpression leads to mitotic defects. **a)** Mitotic progression in 293F cells. Cells were treated with Doxycycline (Dox) for 24 hours, or not treated (Ctrl), and imaged live with SiR-Tubulin. Representative still frames from LSFM videos in control and Dox-treated 293F cells are shown at different time points. Gaussian blur of 0.5 was applied to the images. Black arrows indicate the spindle pole just prior to metaphase onset. Red arrow indicates extra mitotic centriole which disappears at metaphase. Scale bar = 10 *μ*m. Note that in the total time (33 minutes) the Dox-treated cell does not progress beyond metaphase. **b)** Metaphase duration. Average metaphase time of cells which successfully completed mitosis. Quantification was made from FLSM timelapses. Mean ± s.d. from n = 31 ctrl, and 21 Dox-cells, n= 2 independent experiments. Students T-test shows significant difference. **c, d**) Chromosome alignment during metaphase. Cells were treated with Doxycycline (Dox) for 24 hours, or not treated (Ctrl), fixed and stained with antibodies against tubulin (gray). DNA was visualized with DAPI (magenta). In (**c**) representative confocal images are shown. Arrowheads indicate misaligned chromosomes. Scale bar = 5 *μ*m. In (**d**) the percentage of spindles containing mis-aligned chromosomes at metaphase was quantified from. Shown is the mean ± s.d. from 33 control and 38 Dox-spindles, from 3 independent experiments. Students T-test shows significant difference. **e, f**) Spindle length during metaphase. Cells were treated with Doxycycline (Dox) for 24 hours, or not treated (Ctrl), fixed and stained with antibodies against acetylated tubulin (gray). DNA was visualized with DAPI (magenta). In (**e**) representative confocal images are shown. Dotted lines indicate the pole-to-pole distance. Scale bar = 5 *μ*m. In (**f**) spindle length, or pole-to-pole distance was quantified from images such as depicted in (**e**). n= 28 control, and 33 Dox-induced cells from 3 independent experiments.

One of the major activators of the SAC is BUBR1, which together with BUB3 and MAD2 form the Mitotic Checkpoint Complex (MCC) ^36, 39, 40^. The MCC keeps the SAC active until all chromosomes are correctly aligned at the metaphase plate, at which point it is switched off. Consistent with the misalignment phenotype, we found aberrant BUBR1 accumulation at the metaphase plate in Dox-treated cells (Fig. S 2a). Western blotting for BUBR1 in cell lysates did not show a change in protein level after Dox-induction (Fig. S 2b). These data suggest that mild tubulin-overexpression engages the SAC and delays anaphase onset.

Finally, we found that metaphase spindles had an overall reduced length after Dox-addition (Fig. 3e, f). Furthermore, the mitotic defects were accompanied by mis-localisation of spindle-associated proteins (Fig. S 2c-e), as the levels of the kinesin KIF2A, which localizes at spindle poles where it acts to depolymerize MTs ^41–43^, were reduced in Dox-induced cells (Fig. S 2c, d), while CLASP2, which is mainly localised to kinetochore MTs ^44–46^, was increased after Dox-induction (Fig. S 2e). Therefore, spindle formation and architecture are affected in tubulin-overexpressing cells.

Taken together, our results suggest that tubulin overexpression results in altered MT behaviour and mitotic defects. Upon entry into mitosis the tubulin/MT ratio has been shown to suddenly increase ^47^, likely because MTs are rapidly broken down. During mitosis the tubulin/MT ratio is lowered again, and this ratio is subsequently maintained, also upon exit from mitosis and entry into G1 ^48^. We propose that when mitosis begins in Dox-induced 293F cells tubulin levels increase dramatically, since these cells, which already have a surplus of soluble tubulin, also break down their excess MT network. An abnormally high tubulin concentration might, for example, increase the frequency of branched MT nucleation leading to the formation of shorter spindles containing more MTs ^49^. The excess of tubulin during mitosis in Dox-induced cells leads to the activation of the SAC and eventually arrests cells in metaphase.

### Tubulin overexpression deregulates the cell cycle and affects genome integrity

Having established the effects of tubulin overexpression on MT behaviour and mitosis we next explored whether other repercussions could be observed. We first examined the cell cycle in tubulin overexpressing cells using flow cytometry. Cells were fixed after 24 or 48 hours of Dox-induction, and stained with propidium iodide (PI) to investigate DNA content. No effect of Doxycycline was observed in the parental TetR-expressing cell line after 24 hr Dox-induction (Fig. S 3a), indicating that neither Doxycycline itself nor rTTA expression influences the cell cycle. By contrast, we detected a decrease in the G1 population and an increase in the G2/M fraction after 24 hour of tubulin overexpression (Fig. 4a, b). Surprisingly, despite the reduction in the G1 fraction, similar S-phase cell populations were detected in the control and dox-induced cells (Fig. 4a, b). After 48 hr of Dox-induction all fractions (G1, S and G2/M) were perturbed (Fig. 4c). Moreover, DNA content was highly aberrant after 48 hr, with large proportions of cells containing < 2n or > 4n chromosomes (Fig. 4d). Thus, persistent overexpression of tubulin affects the cell cycle and after 48 hr it results in severe DNA abnormalities, suggestive of CIN ^50^.

**Figure 4.**
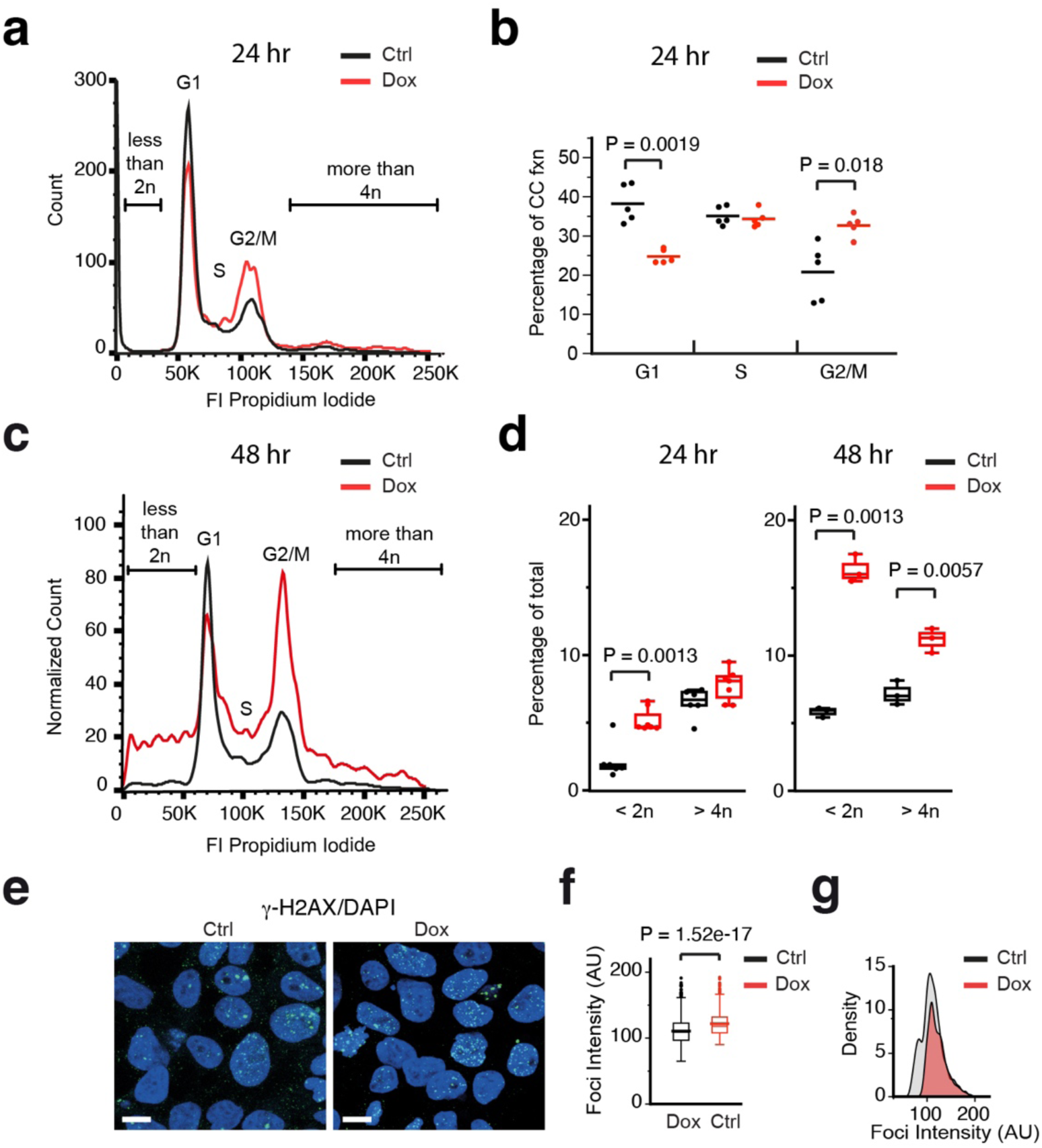
Tubulin overexpression affects the cell cycle and genome integrity. a-d) Cell cycle analysis of 293F cells. Cells were either treated with Doxycycline (Dox, red) or not treated (Ctrl, black). Treatment was either for 24 hr (**a, b**) or for 48 hr (**c, d**). Cells were fixed, stained with propidium iodide, and analysed by flow cytometry. In (**a, c**) representative flow cytometry plots are shown. FI: fluorescence intensity. In (**b, d**) whisker plots of normalised FI are shown, with mean values indicated. n = 6-7 replicates per condition pooled from 3 independent experiments. Student T-Tests were done to reveal significant differences between Ctrl and Dox. **e**) *γ*-H2AX localisation in 293F cells. Cells were treated with Doxycycline (Dox) for 24 hours, or not treated (Ctrl). Cells were fixed, and stained with anti-*γ*-H2AX antibodies (green) and DAPI (blue) to visualise the nucleus. A maximum intensity projection of 7 slices is shown. Scale bar: 10 µm. **f, g**) Quantification of *γ*-H2AX foci intensity. In (**f**) a whisker plot of the foci intensity in arbitrary units (A.U.) is shown, mean values are indicated by wider lines. Student T-Test reveals significant difference between Ctrl and Dox. In (**g**) the foci intensity distribution is shown. Data are from N=2 independent experiments, intensity was measured in more than 1000 foci for each condition.

After mitosis the G1/S checkpoint is the first major cell cycle checkpoint encountered. When satisfied, cells enter S phase to replicate their DNA. G1 release is in part effected by relieving the Cyclin D/Cdk4- and Cyclin E/CDK6-mediated inhibition of the transcriptional complex E2F and Retinoblastoma protein (RB). This occurs, among others, by phosphorylation of RB ^51–53^. Interestingly, the level of phospho-RB was similar in Dox-induced and control 293F cells (Fig. S 3b), despite a decreased number of G1 cells in the Dox-induced population. We therefore hypothesised that tubulin overexpression leads to a premature release of the G1/S restriction point. If true, Dox-induced cells replicate their DNA prematurely, while having a shortage of S phase factors, and cells may experience replication stress, including DNA damage. Using an antibody against *γ*H2A-X, which detects the phosphorylated form of Histone H2A at double strand break (DSB) lesions ^54, 55^, we detected no significant increase in the number of *γ*H2A-X-positive foci observed per nucleus (Fig. S 3c), but did observe a higher intensity of *γ*H2A-X signal within foci after Dox-induction (Fig. 4e-g), suggesting that tubulin overexpression results in an accumulation of DSB lesions due to slower repair.

Combined, our data show that tubulin overexpression affects all three major cell cycle checkpoints, i.e. G1/S, G2/M, and the SAC in mitosis. The premature release of the G1/S checkpoint upon tubulin overexpression results in inadequate levels of DDR and replication factors during S phase, and hence in DDR defects, which, together with excess tubulin itself, may block cells at the G2/M checkpoint. There is evidence for many connections between DDR, MTs and MAPs. For example, MAPs can bind at damaged DNA foci to facilitate repair, interact directly with known DDR proteins, or mediate the transport/localisation of DDR proteins to damaged foci ^56–59^.

Tubulin overexpression in 293F cells might affect such relationships. Conversely, DNA damage is linked to MT stability, as studies in RPE-1 and MCF7 cells have demonstrated that DSBs can specifically induce centrosomal MT-polymerisation in a phenomenon termed DSB-induced MT dynamics stress response ^60^.

### RNA-sequencing reveals autoregulation in tubulin overexpressing cells

To analyse the genome-wide consequences of tubulin overexpression, we performed RNA-sequencing (RNA-seq) on control and Dox-induced cells. Samples were either collected at the start of the experiment (0 hr), or after 24 or 48 hr. Table S1 lists normalised reads of all samples (note that the 0 hr experiment was performed once, in duplicate, while the 24 and 48 hr RNA-Seq experiments were performed twice, once with duplicate samples, and once with triplicates). Principal component analysis (PCA) of the 0 and 24 hr data showed that the 24 hr Dox-induced samples clustered together and were separated from the 24 hr control and the 0 hr samples (Fig. 5a). PCA of the 48 hr samples also showed separation of Dox-induced samples from controls (Fig. S 5a). Altogether, the PCA confirmed the quality of the experiments and suggested that transcriptomic changes do occur in Dox-induced cells.

**Figure 5.**
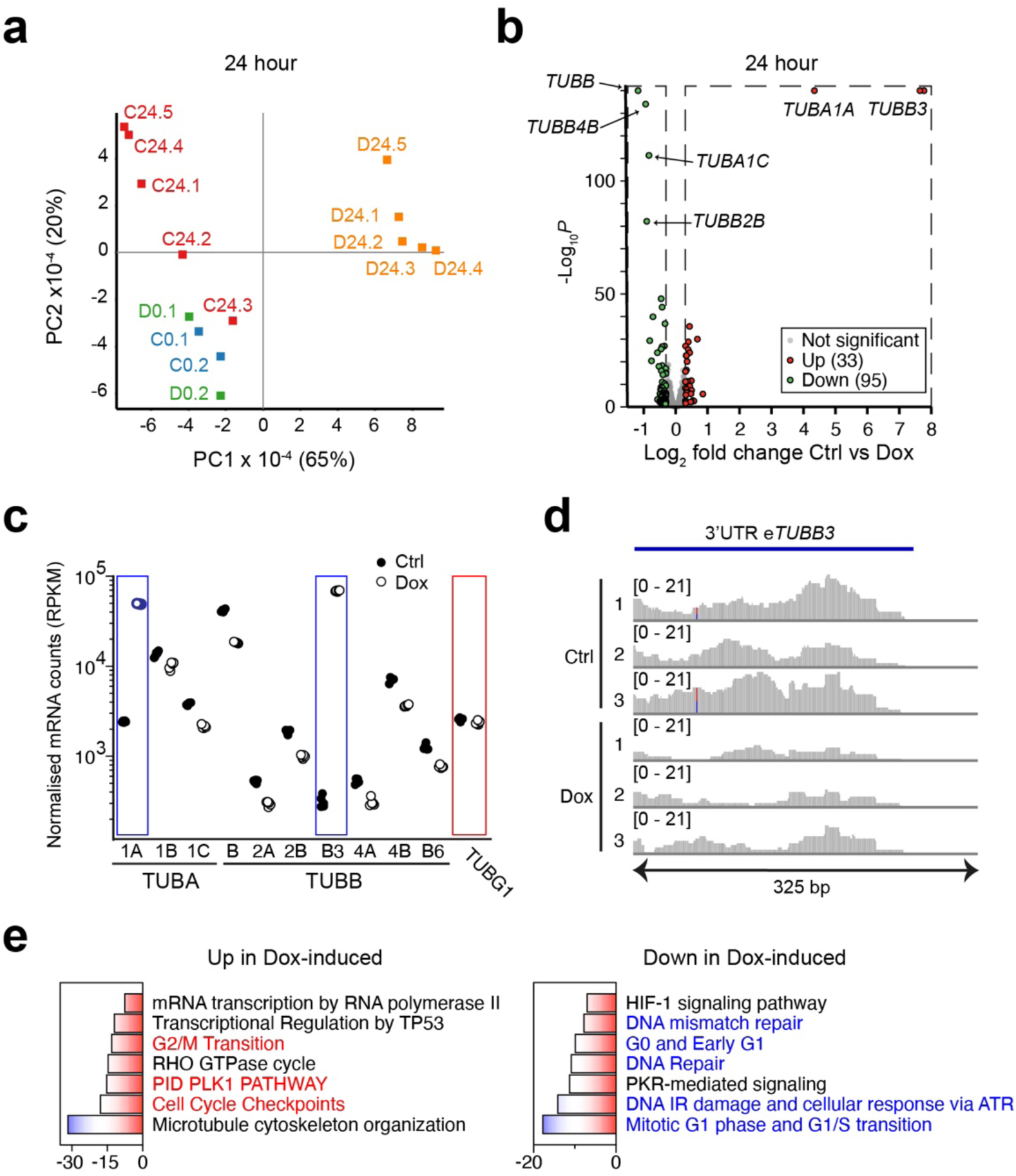
Tubulin overexpression affects the transcriptome. **a)** Principal component analysis of RNA-Seq datasets. Principal component analysis (PCA) was performed on RNA samples derived from control cells (C) or Dox-induced cells (D) cultured for 0 or 24 hr. For the 0 hr timepoint (C0 & D0) we analysed two replicates each for control and Dox-induced cells. For the 24 hr time point two independent experiments were done, in the first we analysed two replicates per line (24.1, 24.2), in the second we analysed three replicates per line (24.3, 24.4 & 24.5). Blue rectangles: control cells, 0 hour, green rectangles: Dox-induced cells, 0 hour, Red rectangles: control cells, 24 hour, orange rectangles: Dox-induced cells, 24 hour. **b)** Volcano plot of differentially expressed genes after 24 hr Doxycycline (Dox)-treatment. Cells were either not treated (Ctrl) or treated with Dox for 24 hr. Green and red points indicate differentially expressed mRNAs (green: down in Dox, red: up in Dox) with p-value < 0.05 and a Log2 fold change of less than -0.3 (green) or more than 0.3 (red). Selected tubulin-encoding mRNAs are indicated. **c)** Tubulin expression in 293F cells. Cells were treated for 24 hours with Dox (open circles), or not treated (Ctrl, black circles). mRNA was isolated and sequenced. Normalised mRNA counts (RPKM) are shown of the indicated mRNAs. Counts for TUBA1A and TUBB3 are highlighted by blue rectangles, note that in the Dox-induced samples they consist of endogenous and recombinant tubulins. Expression of TUBG1 is highlighted with a red rectangle. **d)** TUBB3 3’UTR levels. Cells were treated with Dox for 24 hours, or not treated (Ctrl). mRNA was isolated from the cells and sequenced. The experiment was performed in triplicate. A coverage map of reads on the 3’UTR of TUBB3 was generated in IGV browser. Notice that in Ctrl more reads align to the 3’UTR, indicating that in the Dox-treated cells autoregulation takes place on endogenous TUBB3 mRNA. **e)** Metascape analysis of the set of common deregulated differentially expressed genes after 24 hr Doxycycline (Dox)-treatment. Pathways in red are involved in G2/M transition and mitosis, pathways in blue are involved in G1/S transition and replication stress.

To examine differentially expressed genes (DEGs) we performed a DESeq2 analysis. Not many mRNAs were significantly deregulated after 24 hr Dox-induction (33 mRNAs up and 95 mRNAs down, Fig. 5b, Table S1). By contrast, many more DEGs were observed after 48 hr of Dox-induction (1215 mRNAs up and 1585 down, Fig. S 5b, Table S1), consistent with the more aberrant cell cycle profile seen after 48 hr. Notably, both after 24 hr (Fig. 5b) and 48 hr (Fig. S 5b) tubulin-encoding mRNAs were among the most affected DEGs. To assess tubulin expression in more detail we extracted the reads from tubulin-encoding mRNAs after 24 hr Dox-induction. We observed upregulation of recombinant *TUBA1A* and *TUBB3* in the Dox-induced samples, and downregulation of endogenous tubulin-encoding mRNAs (Fig. 5c, Fig. S 4a). Importantly, the extent of downregulation was similar for all isotypes, strongly suggesting that autoregulation is the mechanism by which tubulin mRNAs are curbed and not transcriptional control, which will not act on all tubulin isotype genes. In addition, although most of the reads from *TUBB3* mapped to the coding region, revealing recombinant tubulin overexpression, we also detected reads mapping to the 3’UTR of *TUBB3*, which showed a reduction in the Dox-induced samples (Fig. 5d). As these reads are derived from endogenous *TUBB3*, the data indicate that autoregulation extends to all endogenous tubulin isotypes.

It has been shown that *TUBG1*, which encodes *γ*-tubulin, a component of the *γ*-tubulin ring complex (*γ*TuRC) ^61^, is co-regulated with tubulin mRNAs in RPE1 cells treated with MTAs ^62^, and that it also is a target of TTC5-mediated autoregulation, despite the fact that its N-terminus is slightly different compared to α / β-tubullin ^63^. Analysis of *TUBG1* reads revealed downregulation in tubulin overexpressing cells, albeit weakly (Fig. 5c). Taken together, our data suggest that tubulin autoregulation is engaged by mild overexpression of the tubulin dimer, and that it persists after 48 hr. As shown above, this system is incapable of restoring normal tubulin levels in the Dox-induced cells. The surplus tubulin is partly converted into MTs, explaining why these are also in excess.

*RNA-sequencing reveals an altered stress response in tubulin overexpressing cells* We next analysed the RNA-seq data for effects beyond autoregulation. We performed both a metascape analysis ^64^ on DEGs (for this analysis we lowered the threshold of the log2 fold expression level change to 0.1, in order to increase the number of DEGs), and a Gene Set Enrichment Analysis (GSEA) on the full RNA-Seq dataset using the hallmark, C2, and C5 gene sets ^65, 66^. Metascape analysis (Fig. 5e, Table S1) revealed downregulation of G1-S- and DNA replication-related pathways in Dox-induced cells (indicated in blue in the figure), and upregulation of G2-related pathways, including PLK1, and of mitotic spindle components (indicated in red in the figure). GSEA confirmed the Metascape results, as the G1-related MCM pathway was down in Dox-induced cells (Fig. S 4b, Table S1). Moreover, GSEA on a locally assembled set of mRNAs encoding cyclins and cyclin-dependent kinases showed that G1/S factors were downregulated in the Dox-induced cells (Fig. S 4c, blue), whereas cyclins and cyclin-dependent kinases important for G2/M were upregulated (Fig. S 4c, red). These results are consistent with our FACS analysis and suggest that shifts in cell cycle fractions partly underlie gene expression differences.

GSEA also revealed enrichment of terms involving +TIPs, MT ends, and motor proteins in Dox-induced samples (Table S1). We extended this analysis by assembling a larger set of +TIPs and performing GSEA on this local dataset. This showed that most of the +TIP-encoding mRNAs, including *CLASP1* and -*2*, were upregulated in Dox-induced cells (Fig. S 4d, Table S1). As tubulin overexpressing cells have an increased G2/M fraction, we hypothesise that in the G2/M phase +TIPs and motor proteins are normally upregulated in 293F cells to support the increased MT dynamicity and MT-based motor protein capacity needed in this cell cycle stage ^67^.

In addition to cell cycle defects, the Metascape analysis suggested that tubulin overexpression affected HIF-1- and PKR-mediated signaling (Fig. 5e, Table S1). Hypoxia-inducible factor 1 (HIF-1) is a transcription factor that regulates the cellular response to oxygen deprivation, or hypoxia ^68^. Cells respond to hypoxia by stabilising the alpha subunits of HIF-1 (HIFA), which subsequently upregulate the transcription of genes required for adaptation to low oxygen conditions ^69^. There are three HIFA proteins (HIF1A, -2A, and -3A). Both HIF1A and HIF3A are expressed in 293F cells, although HIF1A is more abundant (Table S1). Protein kinase double-stranded RNA-dependent pathway (PKR), which is induced by various stress mechanisms including viral infections, acts in the integrated stress response (ISR) and mainly halts protein translation in response to stress by phosphorylating EIF2α, which is involved in translation initiation ^70^. Interestingly, hypoxia can also cause a reduction in translation due to the ISR ^71^, as can amino acid deprivation ^72^, and other types of stress, including mitochondrial dysfunction ^73^. The ISR halts general translation in favour of the synthesis of stress-related proteins that restore protein homeostasis, or proteostasis ^70, 73^. Intuitively, it is logical that stress responses are initiated in cells cultured without a medium refresh, since prolonged culture leads to oxygen deprivation and amino acid starvation ^74^. The fact that HIF-1 and PKR signaling were dampened in Dox-induced cells was surprising and suggested that tubulin overexpression somehow hampers these responses.

### Tubulin overexpression causes mitochondrial and proteostasis defects

Since RNA-Seq does not necessarily detect changes at the protein level, we decided to perform proteomic profiling of Dox-induced 293F cells. We cultured Dox-induced and control cells for 24 hr under 2.5% oxygen conditions (hypoxia) or 20% oxygen conditions (normoxia) and analysed cellular proteomes using mass spectrometry. We detected 8224 proteins in these experiments, of which 3128 passed an ANOVA-based significance threshold (Table S1). Volcano plots of the proteomes in different conditions revealed many more proteins upregulated in normoxia compared to hypoxia, both in control and Dox-induced cells (Fig. 6a, Fig. S 6a). This result is expected since hypoxia can result in general translation inhibition due to the ISR. Surprisingly, we also observed skewing when we compared Dox-induced and control cells, with many more proteins down in the Dox-induced cells, both in hypoxia and normoxia (Fig. 6b, Fig. S 6b). Although the skewing was less pronounced compared to hypoxia versus normoxia (Fig. S 6a,b) the data demonstrate that tubulin overexpression causes proteostasis defects.

**Figure 6.**
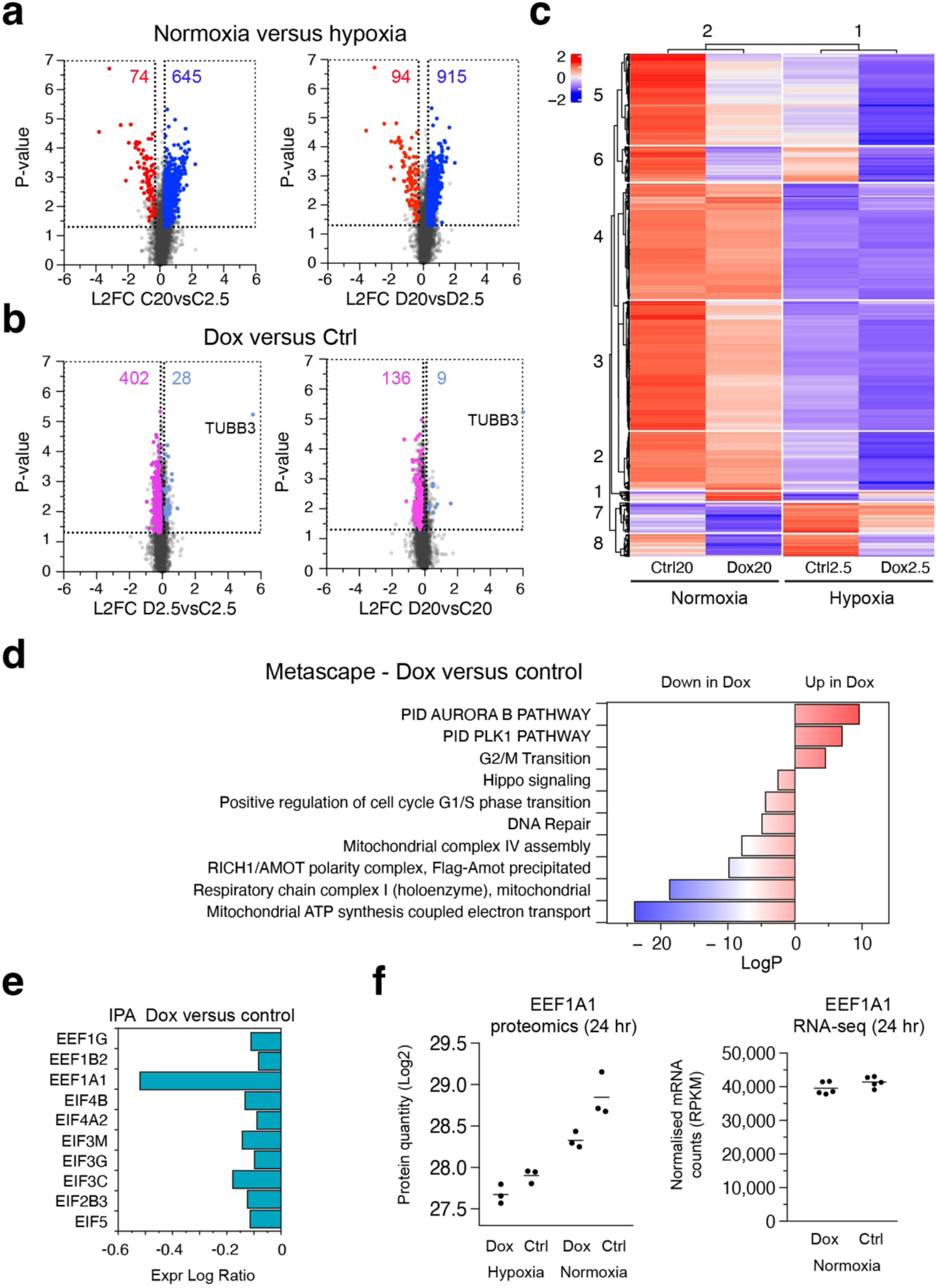
Effects of hypoxia treatment and tubulin overexpression on the proteome. **a**, **b**) Volcano plot of differentially expressed proteins. Cells were cultured for 24 hr in hypoxic (2.5% oxygen) or normoxic (20% oxygen) conditions, and either treated with Doxycycline (Dox) for 24 hr or not treated (Ctrl). Cells were lysed and proteomes were analysed by mass spectrometry. In (**a**) normoxia is compared to hypoxia (left panel: Ctrl cells, right panel: Dox-treated cells). Red dots and numbers indicate proteins significantly downregulated in normoxia,, blue dots and numbers indicate significantly upregulated proteins in normoxia. Qualifying proteins have a log2-fold change (L2FC) of less than -0.3 (red) of more than 0.3 (blue), an ANOVA-based P-value for all samples of less than 0.05, and an individual sample-based significance of less than 0.05. Notice the skewing, i.e. many more proteins are upregulated in normoxia, both in Ctrl and Dox-treated cells. In (**b**) Dox-treated cells are compared to Ctrl (left panel: cells in hypoxia, right panel: cells in normoxia). Pink dots and numbers indicate proteins significantly downregulated in Dox-treated cells, whereas light blue dots and numbers indicate significantly upregulated proteins in Dox-treated cells. The position of rTUBB3, the most upregulated protein in Dox-induced cells, is indicated. Qualifying proteins have a log2-fold change (L2FC) of less than -0.1 (red) of more than 0.1 (blue), an ANOVA-based P-value for all samples of less than 0.05, and an individual sample-based significance of less than 0.05. Notice the skewing, i.e. more proteins are upregulated in Ctrl cells, both in hypoxia and normoxia. **c)** K-means clustering of differentially expressed proteins. We performed k-means clustering on 3228 proteins from the four conditions described in (**a, b**). We divided rows into 8 clusters and columns into two clusters. Scale represents Log2 values of expression. **d)** Metascape analysis of differentially expressed proteins in Dox-induced (Dox) versus control (Ctrl) cells. For visual representation LogP values of proteins upregulated after Dox-induction were inverted to positive values. **e**, **f**) Expression of translation initiation and elongation factors. In (**e**) translation initiation and elongation factors found to be significantly differently expressed (Expr Log Ratio) in an IPA analysis are plotted. The more negative the difference, the less well the factor is expressed in Dox-induced cells (Dox). In (**f**) the left panel depicts protein quantity of EEF1A in the individual samples of Dox-induced (Dox) and control cells (Ctrl), in hypoxia and normoxia, as deduced from the mass spectrometry experiments. The right panel depicts EEF1A mRNA levels in 24 hr Dox-induced (Dox) and control cells (Ctrl) in normoxia, as derived from the RNA-seq experiments.

A heatmap representation of a K-means clustering analysis, where the proteins formed eight groups and the conditions two, visually confirmed the reduced protein levels in hypoxic conditions and after dox-treatment (Fig. 6c). Only in protein cluster 7, which consisted of 195 proteins (Table S1), did we observe increased protein levels in hypoxic conditions (Fig. 6c). This cluster contained the HIF1 response pathway as the top term in a Metascape analysis (Table S1). Thus, both control and Dox-induced cells activate HIF1 in hypoxia. However, HIF1 targets are present at lower levels in Dox-induced cells (Fig. 6c). The HIF1 response is therefore dampened by tubulin overexpression, consistent with the RNA-seq experiments (Fig. 5e). We conclude that tubulin overexpression affects proteostasis and propose that this leads to a weakened response to hypoxia.

To uncover the mechanism by which tubulin overexpression causes proteostasis defects we focussed on the Dox-induced versus control samples. We first analysed differentially expressed proteins (DEPs) using Metascape, and subsequently used Ingenuity Pathway Analysis (IPA) ^75^ on DEPs and an extended protein dataset. Metascape analysis revealed upregulation of terms associated with mitosis and G2/M transition, including the PLK1 pathway, in the Dox-induced cells, whereas G1/S-specific, and DNA repair terms were down (Fig. 6d, Table S1). These data are consistent with our FACS (Fig. 4a, b) and RNA-seq results (Fig. 5e). Surprisingly, a number of terms related to mitochondria were found to be downregulated in the Dox-induced cells, including electron transport, respiration, and mitochondrial complex I and IV assembly (Fig. 6d, Table S1). IPA on DEPs revealed oxidative phosphorylation and respiratory electron transport as the top terms (Table S1), supporting the Metascape results. Since no mitochondrial pathways were detected in the RNA-seq analysis (Table S1), the data suggest that tubulin overexpression affects mitochondrial homeostasis at the protein and not the mRNA level. Moreover, results indicate a specific defect in a subset of mitochondrial proteins related to electron transport. As mitochondrial stress has been shown to lead to proteostasis defects ^73, 76^, we hypothesise that the proteome skewing observed in Dox-induced cells arises due to mitochondrial stress, induced by tubulin overexpression.

Proteostasis defects can be caused by alterations in translation efficiency. To analyse possible effects on translation after tubulin overexpression we performed IPA on an extended protein dataset of 3078 proteins, of which 2819 proteins were down and 259 proteins were up in the Dox-induced cells in normoxia. Strikingly, top hits in IPA revealed terms involving translation, including Major pathway of rRNA processing in the nucleolus and cytosol, Eukaryotic Translation Initiation, Eukaryotic Translation Elongation, and Eukaryotic Translation Termination (Table S1). Many of the proteins in these pathways overlap as they are either mitochondrial or cytoplasmic ribosomal proteins (Table S1). Virtually all proteins in the pathways were downregulated in the Dox-induced cells (Table S1, for visual representation see the Eukaryotic Translation Elongation pathway in Fig. S 6c). Thus, although at the individual level these proteins do not pass the significance threshold for DEPs, the fact that almost all proteins in the different pathways were deregulated in the same manner makes the pathways highly significant. In conclusion, the IPA results suggest that tubulin overexpression slows translation and thereby downregulates many proteins.

Translation is classically divided into three steps, i.e. initiation, elongation, and termination. Transient mitochondrial stress causes translation attenuation due to defects in respiration and ensuing lowered levels of the translation elongation factor EEF1A1 ^76^. On the other hand, prolonged mitochondrial stress eventually results in the ISR and hampers translation initiation ^73^. To distinguish between these two, we examined the levels of significantly deregulated initiation and elongation factors in the Dox-induced proteome, and found that EEF1A1 was most affected (Fig. 6e). Notably, EEF1A1 was downregulated at the protein level in Dox-induced cells, both in normoxia and hypoxia (Fig. 6f, left panel), but not at the mRNA level (Fig. 6f, right panel), indicating that EEF1A1 itself is translationally regulated. Combined, our results suggest that overexpressing tubulin for 24 hr leads to defects in respiration, which results in mitochondrial stress. This, in turn, lowers EEF1A1 causing attenuation of translation and dampened elongation rates. That other initiation and elongation factors are mildly down in Dox-induced cells, as are ribosomal proteins (Fig S6c, Table S1), indicates a mild ISR and general translation inhibition. We propose that tubulin overexpression results in mitochondrial dysfunction leading to stress and attenuated translation elongation. We observe the start of an ISR in these cells, which prevents a proper hypoxic response.

### Characterisation of the tubulome of 293F cells

To uncover tubulin-mediated mechanisms underlying proteostasis defects we investigated the tubulin interactome, or “tubulome”, of 293F cells. We generated a stable 293F cell line in which β-tubulin was tagged at its C-terminus with the ALFA-tag (Fig. S 7a), a small stable α-helix that has been shown to outperform other tags in protein pull down studies ^77^. Using beads coupled to ALFA-tag antibodies, we affinity-purified recombinant tubulin and tubulin-associated proteins (TAPs) from cell lysates of tubulin overexpressing cells, using non-induced cells as controls. In one experiment we washed beads three times (experiment 1) whereas in another we washed beads four times (experiment 2) to distinguish between lower and higher affinity tubulin interactions. We then boiled the beads and identified TAPs in bead supernatants by mass spectrometry (Fig. 7a, Table S1). As expected, more TAPs were detected in the first experiment compared to the second (Fig. 7a, Table S1).

**Figure 7.**
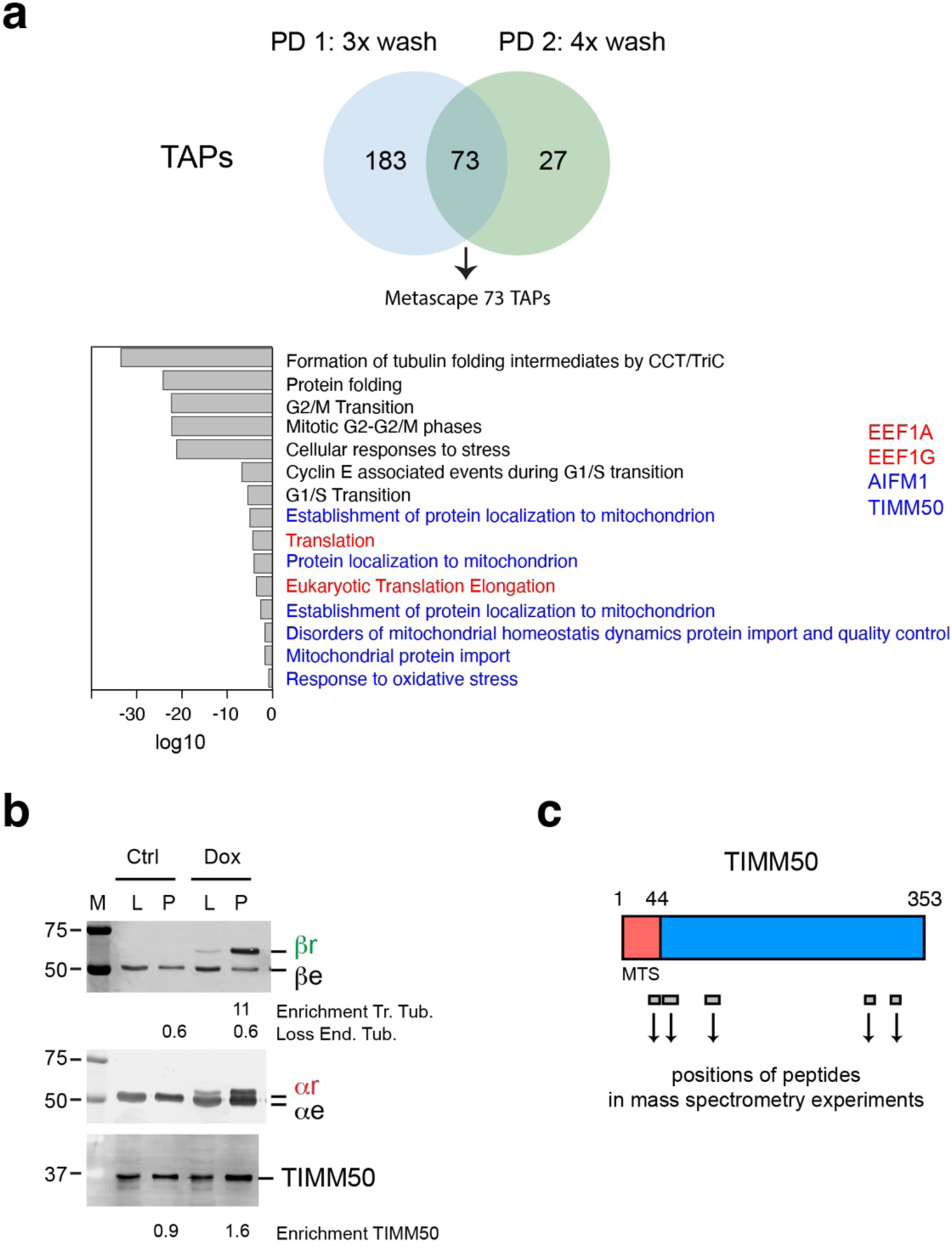
The tubulin interactome. **a)** Analysis of tubulin-associated proteins (TAPs). 293F cells expressing ALFA-tagged tubulin were treated with Dox or not treated. Cells were lysed and recombinant tubulin and TAPs were affinity-purified. In one experiment beads were washed three times (PD 1), in another experiment beads were washed four times (PD 2). Metascape analysis on the 73 TAPs found in both pull downs revealed term associated with process affected by tubulin overexpression (e.g G1/S, G2/M transition) as well as terms associated with mitochondrial localisation, organisation and stress (indicated in blue) and translation (indicated in red). Four proteins (EEF1A1, EEF1G, AIFM1, TIMM50) were related to these terms. **b)** Validation of the tubulin-TIMM50 interaction. Cells were treated as described in (**a**). Tubulin-TAP-containing beads were washed three times after which beads were boiled, and supernatants analysed by western blot using the indicated antibodies (βr, recombinant beta-tubulin, βe, endogenous beta-tubulin, αr, recombinant alpha-tubulin, αe, endogenous alpha-tubulin). Note that TIMM50 is mainly present as ∼35 kDa protein, which represents the precursor form of the protein. The numbers below the blot indicate the relative enrichment of transgenic β-tubulin (Tr. Tub., or βr) and TIMM50 as well as the loss of endogenous tubulin (End. Tub, or βe). The enrichment was calculated by dividing the intensity of the signal in the P lane by the intensity in the L lane. Ctrl: control cells, Dox: Dox-treated cells. L: lysate, P: pull down (bead material), M: marker protein lane. **c)** TIMM50 coverage. The position of the peptides identified by mass spectrometry in the experiments described in (f) is indicated in full length TIMM50. Amino acids 1-44 contain the mitochondrial targeting sequence (MTS) of TIMM50.

We observed 73 TAPs coming down in both pull downs (Fig. 7a, Table S1). Metascape analysis of these common TAPs revealed tubulin folding as the strongest term (Fig. 7a, Table S1). Since chaperones and tubulin folding factors are well known TAPs, these results validate our tubulin interaction approach in stably expressing 293F cells. Importantly, Metascape analysis suggested that common TAPs are involved in the very processes affected by tubulin overexpression in 293F cells. For example, we detected TAPs involved in G2/M and G1/S transition (Fig. 7a, Table S1), and in DNA damage checkpoints (Table S1). In addition, five terms involving mitochondrial homeostasis were observed, including mitochondrial import and quality control (Fig. 7a, indicated in blue, Table S1), and two terms involving translation (Fig. 7a, indicated in red, Table S1). Interestingly, we found the translation elongation factors EEF1A1 and EEF1G as common TAPs (Fig. 7a, indicated in red, Table S1), indicating that elongation factors interact with tubulin. Our tubulin interactome results therefore indicate that tubulin could interfere with mitochondrial homeostasis, G1/S and G2/M transitions, DNA repair, and translation, by binding factors involved in these processes.

The mitochondrial terms consistently contained the proteins AIFM1 and TIMM50 (Fig. 7a, Table S1). Since mitochondrial stress can cause proteostasis defects by attenuating translation ^76^, we followed up on these mitochondrial TAPs. AIFM1 (Apoptosis Inducing Factor Mitochondria Associated 1) is a mitochondrial flavoprotein that was first associated with apoptosis, but was later found to be required for the normal expression of major respiratory chain complexes ^78^. TIMM50 is involved in the import of nuclear-encoded mitochondrial proteins into mitochondria ^79^. We validated the interaction between TIMM50 and recombinant tubulin on western blot using antibodies against TIMM50 and tubulins (Fig. 7b). We note that the relatively weak enrichment of TIMM50 on beads containing recombinant tubulins (Fig. 7b, lower blot, Table S1) as compared to the enrichment of the recombinant tubulins themselves (Fig. 7b, upper and middle blots, Table S1) is well explained, first by a low affinity of the tubulin-TIMM50 interaction itself, and second by competition for TIMM50 between recombinant tubulins, which are enriched after washing, and the much larger reservoir of endogenous tubulins, which are lost (together with TIMM50) after washing.

We found that AIFM1 and TIMM50 are TAPs in our previously published tubulin interactome of HEK293T cells ^13^ . To corroborate interactions further, we performed a tubulin pull down in HeLa cells using the previously published strategy ^13^ (Fig. S 7b). We again observed AIFM1 and TIMM50 (Fig. S 7c, Table S1). Thus, AIFM1 and TIMM50 are consistently identified as TAPs using different purification strategies, tags, and cell lines. TIMM50 is mainly present at the inner mitochondrial membrane, which is not a cellular site where tubulin is expected, leading us to question whether this interaction is physiologically relevant. To reach its target membrane TIMM50 is produced as a precursor protein in the cytosol, with the first 44 amino acids serving as a mitochondrial targeting sequence (MTS). This MTS is cleaved off upon entry into the intermembrane space of mitochondria ^79^. Strikingly, when we mapped the peptides detected by mass spectrometry to TIMM50, we found that one of them was located within the MTS (Fig. 7c). These results suggest that tubulin interacts with the precursor form of TIMM50. In the 293F cell lysates TIMM50 is actually mainly detected in its precursor form (Fig. 7c, see lanes labelled L). Based on our results we propose that excess tubulin binds and sequesters TIMM50, thereby hampering protein import into mitochondria, specifically of proteins involved in electron transport and respiration.

### Regulation of the MT cytoskeleton during cellular stress

We have shown that overexpression of tubulin affects the stress response of cells. We then asked whether the converse also occurs, namely that stress responses affect tubulin levels. We found evidence for this hypothesis in a landmark study which showed that autoregulation is active in many experimental conditions in cells and even *in vivo* ^62^. As part of a dataset verification, the authors examined glutamine (Gln) deprivation in U2OS cells and observed a decrease in tubulin at the mRNA and protein level, as well as a reduction in MTs, after 24 hr deprivation ^62^. A decrease in tubulin at the protein level could, at least partly, be a result of general translation inhibition due to the ISR that sets in upon Gln deprivation. However, a reduction in all of the tubulin-encoding mRNAs is best explained by autoregulation.

At the time the authors only examined one sample and did not discuss a link between stress and autoregulation ^62^. To address this, we retrieved multiple transcriptomic datasets in which either hypoxia or Gln deprivation were studied. We then extracted log2-fold changes in expression of tubulin-encoding mRNAs and selected validated stress-related mRNAs as controls (see Materials & Methods for details). We observed downregulation of virtually all the alpha- and beta-tubulin-encoding mRNAs, both after Gln deprivation and hypoxia (Fig. 8a, b). By contrast, stress-related genes were upregulated, as expected (Fig. 8b). Since the tubulin-encoding mRNAs are regulated in the same manner, the results indicate that autoregulation is activated concomitant with the response to amino acid or oxygen deprivation. Interestingly, *TUBG1* was downregulated together with tubulin-encoding mRNAs (Fig. 8b), as would be expected if autoregulation is activated ^63^.

**Figure 8.**
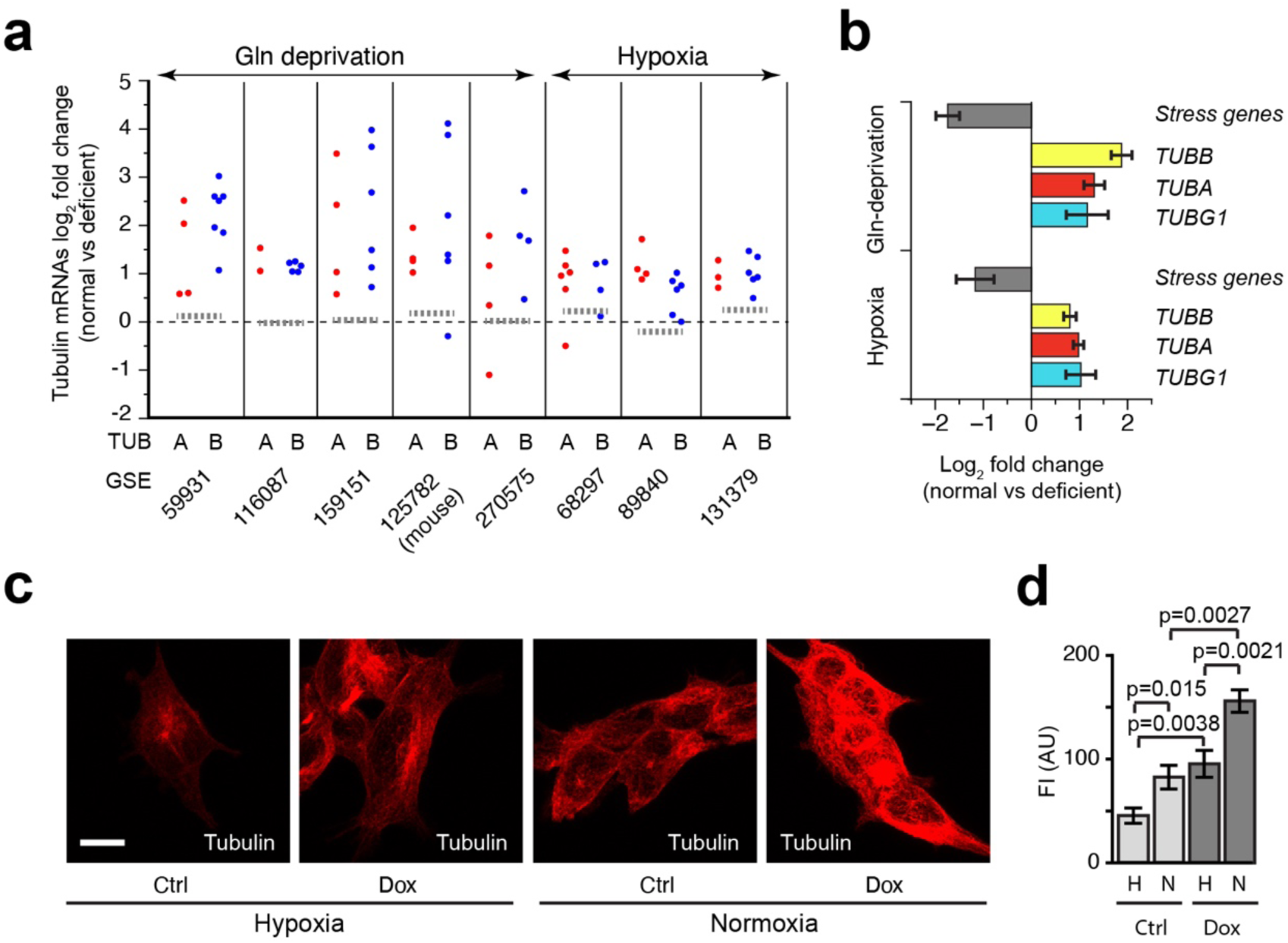
Impaired stress responses in tubulin overexpressing 293F cells. **a)** Tubulin mRNA levels after glutamine-deprivation or hypoxia. We analysed the levels of mRNAs encoding alpha-tubulin isotypes (red dots) or beta-tubulin isotypes (blue dots) in the indicated experiments, where either glutamine (Gln) was deprived (GSE59931, GSE116087, GSE159151, GSE125782, GSE270575) or hypoxia was induced (GSE68297, GSE89840, GSE131379). GSE125782 is from mouse cells. All other datasets are from human cells. **b)** Anti-correlation between stress-induced and cytoskeletal genes. We calculated the log2 fold change in the levels of the indicated mRNAs in normal versus deficient conditions (Gln-deprivation or hypoxia), taking the fold change values of all datasets (Table S1). We selected several known stress genes for glutamine (Gln) deprivation and hypoxia (see Table S1 for the chosen genes). Average log2 fold change is shown ± SEM. **c**, **d**) Effect of hypoxia on the MT network. Cells were cultured for 24 hr in hypoxic (2.5% oxygen) or normoxic (20% oxygen) conditions, and either treated with Doxycycline (Dox) for 24 hr or not treated (Ctrl). Cells were fixed in methanol and stained with antibodies against tubulin. In (**c**) representative images are shown, in (**d**) quantifications of the fluorescence intensity (FI) are shown in arbitrary units (AU). N = 8 Dox-20%, 10 Ctrl-20%, 12 Dox-2.5%, 11 Ctrl-hypoxia 2.5%. T-tests reveal significant differences. Scale bar: 10 micron.

To assess whether stress affected MTs in our system, we cultured control and Dox-induced 293F cells for 24 hr under hypoxic or normoxic conditions, and analysed the network using immunofluorescence microscopy and tubulin antibodies. We found that hypoxia significantly reduced cellular MT density in control cells (Fig. 8c, d), consistent with the reduction in tubulin- and TUBG1-encoding mRNAs seen in the hypoxic RNA-seq datasets (Fig. 8a, b). Since a decrease in MTs has also been observed in glutamine-deprived U2OS cells ^62^, the data suggest that the MT cytoskeleton is dampened in at least two different stress responses. Consistent with the data presented in Fig. 1, the Dox-induced cells displayed a more abundant MT network compared to controls (Fig. 8c, d). Interestingly, hypoxia treatment did decrease the MT network in tubulin overexpressing cells, but not to the extent observed in control cells (Fig. 8c, d). It is possible that an overabundant MT network interferes with the hypoxic response together with surplus tubulin. For example, excess MTs might sequester HIF1A, which has been shown to bind MTs ^80^.

## Discussion

In this work, we utilised a Doxycycline-inducible P2A-based expression system to produce recombinant α/ *β*-tubulin subunits in equimolar amounts. This set-up is controlled in multiple ways. First, no recombinant tubulin is expressed until Dox is added, and by varying the amount of Dox the level of recombinant tubulins can be tuned in an accurate manner. Furthermore, both recombinant α- and *β*-tubulin are expressed, circumventing problems arising from the overexpression of single tubulin subunits. Finally, the construct leaves the autoregulatory MREI motif on recombinant *β*-tubulin free, as well as the C-terminal tail of *α*-tubulin, the site of many PTMs. Here, we generated a stable cell line expressing recombinant TUBB3 and TUBA1A to explore how tubulin overexpression affects cellular homeostasis. By overexpressing a protein that is normally present in cells we create a more natural situation compared to treatment with MTAs, which are xenobiotics.

We show that recombinant tubulins are functional *in vitro* and in cells, and that mild overexpression leads to an excess of soluble tubulin and MTs. The overabundance of MTs indicates that they serve as a storage reservoir for excess tubulin. Although this is logical it does raise the question why, in contrast to tubulin, cells do not sense excess MTs despite the presence of a variety of MAPs, including MT depolymerases. A main reason could be that deviations in MT structure and density are a normal occurrence in cells, serving particular functions. For example, CLASPs locally increase MT density in the periphery of migrating fibroblasts downstream of signaling cues ^81^, among others to control directed motility ^82^. Along these lines it was hypothesised that dynamic MTs act as sensors in cells, and that local deviations in MT density are necessary for mechanical sensing and signaling ^83^. Consistent with this view it was recently shown that mechanical activation rearranges the MT network, leading to the degradation of Angiomotin (AMOT) and altered Hippo signalling ^84^. Notably, we found that the RICH1/AMOT complex and Hippo signaling are downregulated in Dox-induced cells (Fig. 6d, Table S1), indicating that the altered MT network in tubulin overexpressing cells might affect mechanotransduction.

Our RNA-seq analysis showed that a mild overexpression of recombinant tubulins is sufficient to engage autoregulation. Physiological functions of this post-transcriptional regulatory mechanism have not yet been investigated in detail. An early hypothesis was that by controlling tubulin levels autoregulation prevented spurious MT nucleation in cells ^85^. In agreement with this view, we propose that aberrant branched MT nucleation might underlie the mitotic phenotype of Dox-induced cells. Moreover, autoregulation may help to prevent the formation of excess MTs in cells, which seem to lack a MT-based surplus sensing mechanism. A corollary of this is seen in stress situations like Gln-deprivation and hypoxia, where transcriptomic dataset analysis suggests that autoregulation is activated and that this serves to downregulate MTs.

Indeed, immunofluorescence microscopy studies show that this correlates with a diminished MT network in hypoxic 293F cells, and, as shown earlier, with a reduction in MTs in Gln-deprived U2OS cells ^62^. Autoregulation therefore appears to contribute to the downregulation of the MT network in stress situations. It is conceivable that this establishes a specific cellular organization required for a proper stress response. It will be interesting to investigate whether PTX, a major anti-cancer agent which has effects beyond mitosis ^86–88^, kills non-dividing cancer cells because of a failing response to hypoxia or amino acid starvation, or to other types of stress that require a reduction in MTs to reestablish homeostasis. If true, this could be further explored in cancer therapy designs.

We found that tubulin overexpression induces mitochondrial dysfunction. We propose that excess tubulin sequesters TIMM50 and AIFM1 resulting in reduced mitochondrial import. Indeed, constituents of the mitochondrial electron transport chain and oxidative phosphorylation, shown to be transported by TIMM50 ^79^, were downregulated in tubulin overexpressing cells at the protein but not the mRNA level. Our mass spectrometry-based experiments furthermore show that tubulin overexpression results in proteostasis defects and suggest that this occurs by translation attenuation. Whereas the ISR inhibits general translation by phosphorylating EIF2α, acting on translation initiation ^70^, transient mitochondrial stress affects elongation by lowering the level of EEF1A1 ^76^. Strikingly, we found that EEF1A1 protein levels are down in tubulin overexpressing cells. We therefore hypothesise that excess tubulin sequesters AIFM1 and TIMM50, resulting in a lowered mitochondrial protein import and stress, and that this in turn reduces EEF1A1, attenuates translation, and skews the proteome in Dox-induced cells. As we observed an interaction between tubulin and two translation elongation factors (EEF1A1 and EEF1G) in our ALFA-tag pull downs, we presume that tubulin overexpression can directly influence translation elongation. We detected an interaction between tubulin and EEF1G in a previous TAP identification ^14^, showing that this interaction is observed in another cell line using a different pull down approach. In addition, plant elongation factor-1α (EF-1α) was previously found as a TAP in an affinity-based large-scale identification of tubulin-binding proteins using an Arabidopsis cell extract ^89^, indicating the tubulin-EEF1A1 interaction is also conserved.

TTC5 simultaneously binds the nascent translating tubulin polypeptide and the ribosome itself ^10^, and then recruits SCAPER and the CCR4-NOT deadenylase to degrade tubulin mRNAs ^11^. Studies with protein synthesis inhibitors revealed that TTC5 depends on ongoing translation to initiate autoregulation ^90, 91^. For example, cycloheximide at concentrations which block translation elongation and hence immobilise tubulin nascent chains, increases tubulin mRNA degradation ^91^. This is consistent with the notion of a time window for the TTC5-nascent tubulin interaction, which is increased when elongation is blocked. By contrast, inhibition of translation initiation by the addition of pactamycin blocks tubulin mRNA degradation ^91^, in line with the notion that the tubulin nascent chain is crucial for TTC5-mediated autoregulation. In normal cells, translation is rapid and TTC5 therefore has a short time window to interact with the first amino acids of the tubulin peptide before they fold up. This explains, at least partly, why TTC5 activity is low in normal conditions ^12^. We provide evidence here that surplus tubulin slows general translation, increasing the time window for a TTC5-nascent tubulin interaction. TTC5 in turn relays specificity for tubulin mRNA degradation ^10, 11^, and, as recently discovered, for *TUBG1* mRNA degradation ^63^. Whether more mRNAs, encoding cytoskeletal or other proteins, are controlled by TTC5 remains to be shown. Excess tubulin may lower translation elongation both indirectly, via mitochondrial stress, and directly, via interaction with elongation factors. It will be interesting to further explore a correlation between translation attenuation and autoregulation, also in view of our findings that hypoxia and Gln deprivation, which lead to an ISR, activate autoregulation. The latter result, which is seen in eight transcriptomic datasets, would not be expected if translation inhibition were the sole outcome of these stress responses, since translation inhibition blocks autoregulation ^91^.

Sequestering of MAPs and TAPs is an overarching explanation for multiple phenomena observed in this report, with mitochondrial stress serving as a prime example. The premature G1/S release and ensuing replication stress observed in tubulin overexpressing cells could also be caused by the sequestering of an important protein involved in the G1/S checkpoint. Furthermore, as explained above, HIF1 might be sequestered by surplus MTs in Dox-induced cells, preventing a proper hypoxia response. Our RNA-seq data also reveal upregulation of targets of the RHO GTPase cycle (Fig. 5e). This might be due to the titration of the GTP exchange factor GEF-H1 ^83^, which is sequestered in its inactive state on MTs ^92^. A final example is the deregulation of the PLK1 pathway, which we observed in our RNA-seq (Fig. 5e), proteomics (Fig. 6d), and TAP (Table S1) datasets. PLK1 was shown to bind MTs ^93^, and both an excess of tubulin and of MTs could affect PLK1 and thereby influence mitosis. Altogether our data suggest that tubulin has multiple functions beyond the assembly of MTs. Here, we reveal a role in the three main cell cycle checkpoints, in cellular stress responses, and in translation. Our data explain why tubulin, a major cytoskeletal protein and among the more abundantly expressed proteins in a cell, needs to be exquisitely controlled to maintain a healthy genome and proteome.

## Supporting information

Revision plan

## Acknowledgements

This work was supported by grants from the Netherlands Organisation for Scientific Research (TNW 15511).

## Data Availability

For data analysis we downloaded the GSE59931, GSE116087, GSE159151, GSE125782, GSE270575, GSE68297, GSE89840, and GSE131379 datasets from the GEO data repository. RNA-sequencing datasets generated in this study will be made available in the GEO data repository upon acceptance of this manuscript.

## Methods

### Reagents, antibodies, and plasmids

We used the following non-standard reagents: Hygromycin B (50 mg/mL, Invitrogen, 10687-010), Puromycin (Sigma, P7255), Doxycycline hyclate (Sigma, D9891), DMEM (Gibco), Penicillin/Streptomycin (Gibco), Gibson Assembly Master Mix (NEB, E2611L), Protease Inhibitor Tablets EDTA-free (Roche Diagnostics, 11873580001), Ni-NTA Agarose beads (Qiagen, 30210), Ni-NTA Superflow Cartridge (1 mL) (Qiagen, 30721), Imidazole (Sigma, D1916), Sir-Tubulin (Spirochrome, sc002), Propidium Iodide (Invitrogen, P3566), Prolong Gold with DAPI (Life Technologies, P36931).

We used the following antibodies: anti-CLASP1 (Absea, KT66), anti-alpha-tubulin (Abcam, ab18251 (YL1/2; rat) and ab7291 (DM1A; mouse)), anti-beta-tubulin (Sigma T8328, mouse), anti-acetylated tubulin (Sigma T6793, mouse), anti-detyrosinated tubulin (Abcam, ab48389, rabbit), anti-KIF2A (Novus Biologicals, NB500-180, rabbit), anti-GAPDH (Absea, KT186, mouse), anti-Phospho-Rb (Ser807/811, CST, 8516), anti-Rb (CST, 9309), anti-BubR1 (Bethyl Labs, A300-995A-T). We used the following secondary antibodies: Goat anti-Rabbit IgG, Alexa Fluor 594 (Thermo Fisher, A-11037), Goat anti-Rabbit IgG, Alexa Fluor 488 (Thermo Fisher, A-11034), Goat anti-mouse IgG, Alexa Fluor 594 (Thermo Fisher, A-11005), Goat anti-mouse IgG, Alexa Fluor 488 (Thermo Fisher, A-11001), Goat anti-Rat IgG, Alexa Fluor 594 (Thermo Fisher, A-11007), Goat anti-Rat IgG, Alexa Fluor 488 (Thermo Fisher, A-11006), IRDye, 680RD Goat-anti Rat IgG (Licor, 926-68076), IRDye, 680RD Goat-anti Rabbit IgG (Licor, 926-68071), IRDye, 680RD Goat-anti Mouse IgG (Licor, 926-68070), IRDye, 800CW Goat-anti Rabbit IgG (Licor, 926-32211), IRDye, 800CW Goat-anti Mouse IgG (Licor, 926-32210).

For cell line generation and transient transfections various plasmids were used and constructs generated. rTTA-N144 (Addgene, 66810) expresses the reverse Tet transactivator (rTTA) under the EF1a promoter and the hygromycin resistance gene under the PGK promoter. The pTET-TRE vector (originally from Clontech, 631167) was a kind gift from Robbert Rottier (Department of Pedriatic Surgery, Erasmus MC). The TUBA1A and TUBB3 cDNAs were amplified from an existing pcDNA3-based dual plasmid ^13^. The TEV, 6xHIS, and P2A sequences were synthesized either as primers or as short gene blocks (IDT). Sequences were assembled together using Gibson Assembly. Constructs were then used to create other constructs, for example with an ALFA tag inserted after TUBB3. The bio-Sumo*-GFP-based dual tubulin constructs have been described ^13^, as has the EB3-GFP plasmid ^21^.

### Standard molecular biology methods

For RNA isolation, cells were collected and resuspended in 1 ml TRIzol (Sigma) and incubated for 5 min at 30 °C. 200 μl phenol-chloroform (Sigma) was added and samples were incubated on a shaker for 3 min at 30 °C. Samples were subsequently centrifuged for 15 min, 12.000 rpm at 4 °C in a table-top centrifuge. The aqueous phase (top layer) was collected and 250 μl 100% ethanol (Sigma) was added. Samples were then transferred to RNeasy spin columns (Qiagen) and processed according to the manufacturer’s instructions.

SDS-PAGE was carried out according to standard procedures using the mini-PROTEAN system (Biorad). After electrophoresis gels were either fixed and stained with Coomassie Brilliant Blue R-250 or blotted onto a PVDF (Millipore) membrane through wet-transfer for 2 hours at 4°C. The membrane was then blocked with 5% skim milk (Sigma) in phosphate-buffered saline (PBS) and 0.1% Tween-20 (PBS-T), for 30 minutes, and incubated overnight (O/N) at 4°C with primary antibody. Membranes were washed 3 times with PBS-T and incubated for 45 minutes with secondary antibody. After 3 washes with PBS-T, membranes were imaged using the Odyssey CLx (Licor).

### Stable cell lines and transient transfections

To generate stable cell lines expressing dual tubulin under control of the reverse-Tet transactivator (rTTA) protein ^15^ the suspension cell line 293F (Life Technologies, R790-07) was passaged in 125 mL spinner flasks (Corning) in serum-free medium (Freestyle, F17, Gibco) supplemented with L-Glutamine. Cells were moved to plastic cell culture dishes (10 or 15 cm) for adherent culturing in DMEM (Gibco) supplemented with 10% FCS and 1% P/S (Gibco) for 2-3 passages in DMEM supplemented with 10% FCS and 1% P/S. To generate a stable rTTA-expressing line, cells were transfected with rTTA-N144 (Addgene, 66810) using XtremeGene-HD reagent (Roche). Two days later, Hygromycin (Sigma) was added at 100 *μ*g/ml, and cells were cultured for at least 10 days to allow resistant clones to form. 24 clones were picked and screened by transient co-transfection with a tubulin expression construct followed by western blotting. A positive rTTA-expressing clone was selected for the next rounds of transfection.

We generated two stable cell lines with pTRE-dual expression constructs, which are depicted in Fig. 1a (His-based) and Fig. S 7a (ALFA-based), respectively. Clones were selected in Puromycin (1 *μ*g/ml), and screened via western blotting for alpha tubulin for the presence of recombinant tubulin after DOX addition.

For tubulin interactome analysis we transiently transfected HeLa cells with dual tubulin constructs described previously, i.e. Bio-Sumo*-TUBB3-P2A-GFP-TUBA1A and TUBB3-P2A-Bio-Sumo*-GFP-TUBA1A ^13^. To visualise MT growth in tubulin overexpressing cells, 293F cells were seeded on 24 mm coverslips in 6-well plates. The day after, cells were transfected with 1 µg EB3-GFP plasmid per 6-well using Lipofectamine 2000 according to the manufacturer’s instructions. Doxycycline was added to the cells 4 hours post-transfection and imaging of cells was performed 24 hours later.

### Cell cycle analysis

Cell Cycle analysis was performed using Propidium Iodide (PI). 1x10^6^ cells were harvested and centrifuged in PBS at 500xg, followed by resuspension in 800 *μ*L ice-cold PBS. Then ice-cold ethanol was added dropwise to a final concentration of 66% to fix the cells. Cells were incubated for 90 minutes in the cold, then slowly equilibrated to room temperature. Cells were then washed once with PBS containing 1% BSA, then resuspended in 500 mL of PI staining solution containing 50 *μ*g/ml RNase A and 50 *μ*g/ml PI in PBS/BSA. Cells were analysed immediately on a Flow Cytometer. For data analysis, FlowJo v.10.8 and the built-in cell-cycle quantification tool were used (the univariate model without any adjustments).

### Tubulin purification from 293F cells

For protein purification, cells were seeded at 1x10^6^/mL in suspension culture and grown in autoclaved glass spinner bottles fitted with magnetic stirrers in the method previously described ^94^ and maintained with constant stirring at 37^0^C and 5% CO_2_ in a humidified incubator. 500 mL of rTTA-293F cells cultured were treated with 5 *μ*g/mL Doxycycline for 3 days. Cells were harvested and dounced in a dounce homogenizer in lysis buffer (100 mM PIPES pH 6.8, 10 mM MgCl_2_, 5 mM DTT, 300 mM NaCl, 1 mM ATP, 1mM GTP, 3U/mL Benzonase, and Protease Inhibitor Cocktail). The amount of lysis buffer was kept equal to the wet volume of pelleted cells. This was followed by centrifugation at 4° C for 30 min at 55,000 rpm using the 70.1 Ti rotor on a Beckman Optima XE-90 ultracentrifuge. The clarified lysate was carefully removed, supplemented with 10% glycerol and bound to 200 *μ*L Ni-NTA beads for 2 hours at 4° C. Beads were washed 6 times in wash buffer (100 mM PIPES ph 6.8, 10 mM MgCl_2_, 10% Glycerol, 500 mM NaCl, 2 mM EGTA, 0.04% NP-40, 1 mM ATP, 1mM GTP and 50 mM Imidazole. and then eluted in two successive rounds with wash buffer now containing 500 mM imidazole at 4° C. The resulting eluate was pooled and subjected to one cycle of polymerization-depolymerization before being resuspended in MRB80 buffer.

### MT pelleting assays

For the *in vitro* MT pelleting assay 20-40 *μ*L of purified tubulin eluate in MRB 80 buffer, obtained as described above, was supplemented with an additional 10% glycerol, then incubated at 37^0^C with or without 10 *μ*M Paclitaxel. A similar reaction was incubated at 4^0^C without Paclitaxel as negative control. After 1 hour, samples were centrifuged on a Beckman airfuge at room temperature, for 15-20 minutes at 100,000g. The supernatant, and pellet were collected separately, and boiled in SDS-PAGE gel loading buffer such that the final loading volume was identical for both conditions.

The *in vivo* MT pelleting assay was performed as described previously ^95^. Briefly, cells were grown on 10 cm dishes, and Doxycycline (2 *μ*g/ml) was added one day before harvesting. Cells were harvested at ∼70-80% confluency. Tubulin Extraction Buffer (100mM PIPES ph 6.8, 1mM MgCl_2_, 1mM EGTA, 3mM GTP, 30% Glycerol, 5% DMSO, and 1 protease inhibitor tablet per 10 mL buffer) was added to cells at 22^0^C, cells were scraped and centrifuged in a table-top Eppendorf microfuge at 15000 rpm for 25 minutes at room temperature. The Supernatant and Pellet fractions were collected, and boiled with 2X SDS sample buffer. Equal volumes of each fraction were loaded for SDS -PAGE and western blotting. To control for the total amount of cells, GAPDH was used as an internal loading control. Quantification of western blots was performed using the Gel Analysis tool in Fiji.

### RNA sequencing and transcriptomic analysis

RNA-seq samples, prepared as described above, were further processed with an Illumina TruSeq Stranded mRNA Library Prep kit. The resulting DNA libraries were sequenced according to the Illumina TruSeq Rapid v2 protocol on an Illumina HiSeq2500 sequencer, generating single-end reads of 50 bp. Illumina adapter and poly-A sequences were removed from the reads and the trimmed reads were then aligned to the mouse genome using HISAT2 ^96^. After this SAMtools ^97^ was used to sort and merge the alignments from multiple flow cells. We performed quantification using HT-seq count ^98^, after first sorting alignments on readname using SAMTools. HT-seq counts were used as input for DESeq2 ^99^.

As an alternative to the HT-seq approach, and for other analyses, we used Bam files as input for SeqMonk, a tool for viewing and analysing mapped sequence data (https://www.bioinformatics.babraham.ac.uk/projects/seqmonk), followed by principal component analysis (PCA), DESeq2 ^99^, Metascape ^64^, or Gene Set Enrichment Analysis (GSEA) ^66, 100^. For GSEA we also assembled local datasets. Table S1 lists the genes present in these local datasets.

To retrieve transcriptomic datasets in which cells were either deprived of glutamine or cultured in hypoxia, we used iDEP ^101^ (search term “glutamine” or “hypoxia”) or GEO Omnibus (search term “glutamine AND deprivation AND RNA-Seq”). For glutamine deprivation the following datasets were selected: GSE59931, GSE116087, GSE159151, GSE125782, GSE270575. For hypoxia the following datasets were selected: GSE68297, GSE89840, GSE131379. Except for GSE59931, which was a microarray-based analysis, all datasets were based on RNA-seq. All datasets were of human origin with the exception of GSE125782, which was from mouse cells. When possible, we used the authors’ analysis, or we employed GEO2R to perform a differential gene expression analysis, and downloaded tables with Log2-fold changes in gene expression. Alternatively, we downloaded tables with RPKM counts, calculated average gene expression values for the two groups and then derived Log2-fold changes in gene expression. We retrieved and plotted values from all the datasets for α/ *β*-tubulin mRNAs, for *TUBG1*, and for selected mRNAs encoding stress genes (*ATF3, ATF4, ATF5, ATF6, DDIT3, DDIT4, EIF2S3, EIF4EBP1, VLDLR*).

### Mass spectrometry and proteomic analysis

For total proteome analysis 2 X 15 cm dishes of Dox-inducible 293F cells were treated with or without Doxycycline to induce dual tubulin expression for 24 hours. Cells were harvested at 80% confluency, and the cell pellet was flash frozen in liquid nitrogen prior to further handling. 3 replicates per condition were taken for the experiment. Protein samples were extracted and digested with trypsin to generate peptides for mass spectrometric analysis. Peptides were analysed using a data-independent acquisition (DIA) workflow, in which precursor ions within predefined mass-to-charge (m/z) windows are fragmented systematically, enabling comprehensive sampling of peptide ions. Peptide fragmentation was performed using higher-energy collisional dissociation (HCD). Fragmentation occurs primarily along peptide bonds, generating fragment ion series that enable peptide sequence identification. Experimental spectra were matched against predicted spectra derived from protein sequence databases, with machine-learning–based prediction of fragment ion intensities improving identification performance for complex DIA datasets. Peptide abundance was quantified by integrating fragment ion intensities across chromatographic elution profiles. These signal intensities provide a linear estimate of peptide concentration in the digested sample. Peptide identification and protein inference were performed using DIA-NN. Protein inference was based on a maximum parsimony approach that determines the minimal set of protein groups required to explain all observed peptides. Proteins sharing identical peptide evidence were grouped together to account for sequence overlap among isoforms or homologous proteins. Protein annotations were derived from isoform-specific entries in UniPro**t**. Each sample was analyzed twice using different compensation voltage settings in a High-Field Asymmetric Waveform Ion Mobility Spectrometry (FAIMS) interface coupled to the mass spectrometer. This gas-phase fractionation reduces spectral complexity and increases peptide identification depth. Peptide intensities from both runs were combined prior to protein quantification. Protein abundance was calculated using the MaxLFQ algorithm, which estimates protein intensities by integrating pairwise peptide ratios across samples to minimize biases arising from peptide-specific ionization differences. Statistical analyses were performed on log2-transformed protein intensities. Differences between experimental conditions were evaluated using one-way Welch’s ANOVA, which does not assume equal variance between groups. Pairwise comparisons were conducted using the Games–Howell post-hoc test. P-values were interpreted in conjunction with the overall ANOVA result to determine significant differences between conditions.

The mass spectrometry experiment yielded 8224 proteins, of which 3128 passed the ANOVA threshold P-value of 0.05. We generated Volcano plots of differentially expressed proteins (DEPs), comparing normoxia to hypoxia and control to Dox-induced cells. We used a cut-off of 0.3 or -0.3 for significance. DEPs were further analysed by Metascape to document the pathways and processes affected by hypoxia and Dox-induction. We used Complex Heatmap ^102, 103^ to visualise protein expression.

To identify tubulin associated proteins (TAPs) we performed several experiments, as described above. For ALFA pull downs we used 293F cells stably expressing a dual tubulin construct in which the ALFA-tag was coupled to the C-terminus of beta-tubulin (see Fig. S 7a for schematic representation of the dual construct). Cells were grown in 2X15cm dishes per condition and treated for 48 hr with Dox or not treated. After 48 hr medium was refreshed and cells were grown for another 24 hr. Cells were lysed in 80 mM pipes, 300mM KCl, 10% Glycerol, Protease Inhibitor Cocktail (Roche, 1 tablet per 10 ml buffer), 1mM EGTA, 10mM MgCl_2_, 5mM DTT, 1 mM GTP, 0.2% Triton-X-100 (Sigma), 3U/mL Benzonase (EMD Millipore). Recombinant tubulin and TAPs were purified using 50 *μ*l of ALFA bead slurry (ALFA Selector CE, Nanotag Technologies) and washed in 80 mM PIPES, 300mM KCl, 10% Glycerol, 1mM EGTA, 10mM MgCl_2_, 1mM DTT, 1 mM GTP, 0.03% Triton-X-100. In one experiment beads were washed three times (PD1), in another experiment beads were washed four times (PD 2). For biotin-streptavidin-based pull downs (see Fig. S 7b for schematic representation of the dual construct) we transiently transfected HeLa cells and harvested cells using a previously published protocol ^13^.

For ALFA-tag based mass spectrometry SDS-PAGE gel lanes were cut into 1-mm slices and subjected to in-gel reduction with dithiothreitol, alkylation with 2-chloroacetamide and digested with trypsin (sequencing grade; Promega), as described previously ^104^. Nanoflow liquid chromatography tandem mass spectrometry (nLC-MS/MS) was performed on an EASY-nLC coupled to an Orbitrap Fusion Lumos Tribid mass spectrometer (ThermoFisher), operating in positive mode. Peptides were separated on a ReproSil-C18 reversed-phase column (Dr Maisch; 15 cm × 50 μm) using a linear gradient of 0–80% acetonitrile (in 0.1% formic acid) during 120 min at a rate of 200 nl/min. The elution was directly sprayed into the electrospray ionization (ESI) source of the mass spectrometer. Spectra were acquired in continuum mode; fragmentation of the peptides was performed in data-dependent mode by HCD.

Raw mass spectrometry data were analyzed with the MaxQuant software suite ^105^ (version 2.0.1.0) as described previously ^104^ with the additional options ‘LFQ’ and ‘iBAQ’ selected. The false discovery rate of 0.01 for proteins and peptides and a minimum peptide length of 7 amino acids were set. The Andromeda search engine was used to search the MS/MS spectra against the Uniprot database (taxonomy: *Homo sapiens*, release June 2021) concatenated with the reversed versions of all sequences. A maximum of two missed cleavages was allowed. The peptide tolerance was set to 10 ppm and the fragment ion tolerance was set to 0.6 Da for HCD spectra. The enzyme specificity was set to trypsin and cysteine carbamidomethylation was set as a fixed modification. Both the PSM and protein FDR were set to 0.01. In case the identified peptides of two proteins were the same or the identified peptides of one protein included all peptides of another protein, these proteins were combined by MaxQuant and reported as one protein group. Before further statistical analysis, known contaminants and reverse hits were removed. For the ALFA-tag pulldowns, of the proteins identified by mass spectrometry the ones with an IBAQ score >15 were included in the list of TAPs. For the Streptavidin pulldowns, proteins with an IBAQ score >20 were included in the list of TAPs.

For biotin-streptavidin-based mass spectrometry the experiments and analysis were performed as described previously ^13^, using 120 minute LCMS runs, and the MaxQuant and fasta versions described above.

### Immunofluorescence microscopy

Cells cultured on 24 or 18-mm glass coverslips were fixed in either 4 % PFA or -20^0^C methanol depending on the antibody used. After fixation cells were permeabilized with 0.15% Triton X-100 for 10 min, and blocked in PBS containing 0.05% Tween-20 and 1% BSA before incubating in primary antibody diluted in PBS containing 0.05% Tween-20. After incubation for 1 hour at room temperature, coverslips were rinsed three times in PBS containing 0.05% Tween-20 and then incubated with secondary antibodies for 45 min in the dark at room temperature. Coverslips were finally mounted in Prolong Gold containing DAPI (Invitrogen). Pre-extraction of cells was performed by incubating coverslips with PHEM buffer (60mM PIPES pH 6.8, 25mM HEPES, 10mM EGTA, 1mM MgCl_2_, 0.01% TritonX-100 for 1 min at room temperature.

### Image acquisition, processing, and quantification

Fluorescence images of fixed cells were acquired on a Leica SP8 confocal fitted with a DMi8 microscope, using a 63x HCX PL APO CS2 oil objective (1.4 N.A.) with band pass filters set to 415-485 nm for the 405 nm laser, 500-575 nm for the 488 nm laser, and 591-660 nm for the 561 nm laser. All images were acquired at a resolution of 1024 x1024 pixels, and step size for z-stacks was kept at 300 nm unless otherwise specified. For multi-color imaging, frame and line averages were specified individually for each channel as required.

Live cell imaging experiments to measure EB3-GFP-labelled plus-ends were carried out on a Nikon Eclipse Ti-E microscope fitted with a Spinning Disc unit (CSU-X1, Yokogawa), using a Nikon 100x CFI APO TIRF 1.49 N.A. oil objective. The microscope was equipped with a Nikon perfect focus system and QuantEM 512SC EMCCD camera (Photometrics), with an HQ525/50m emission filter (Chroma), and 405, 491, 561 diode lasers (50mW, Cobolt). Images were projected onto the CCD chip of the camera at a magnification of 67 nm/pixel. A stage top heater (INUG2E-ZILCS, Tokai Hot) and lens heater were used to maintain cells at 37°C during imaging. Stream Acquisition was performed using Metamorph 7.5 (Molecular Devices) at a constant frame rate of 2 frames per second. Focus was maintained throughout imaging using the perfect focus system.

For light sheet fluorescence microcopy (LSFM) cells were seeded onto Ibidi TruLive imaging chambers one day prior to imaging. The chambers themselves were cleaned with a plasma cleaner for 3 min prior to seeding. On the day of imaging, the cells were incubated with 1 µM Sir-Tubulin dye for 3 hours followed by washout and incubation in fresh pre-warmed medium. The Ibidi TruLive imaging chambers were then placed in an InviSPIM light-sheet microscope (Luxendo, Bruker) equipped with a Nikon CFI Plan Fluor 10x/0.3 NA illumination water objective, a Nikon CFI Apo 25x/1.1 NA detection water objective, and 2 high-speed sCMOS cameras (Hamamatsu Orca Flash 4.0 V3 2048 × 2048 pixels). For imaging, we used the 633 nm laser set to 15% and the 656 long pass filter. Acquisition was performed using the LineScan mode set at 25 pixels width. 3D images were acquired with the bessel light sheet with a thickness of 35, and 400 ms exposure time. The z-step size was 1 micron. Images were acquired every 3 min for up to 3 hours. Maximum intensity projections (MIPs) of the z-stacks were taken for analysis of metaphase duration.

Measurement of *γ*-H2AX was performed in cells fixed in 4% PFA and stained with anti-*γ*-H2AX antibodies. Z-stacks were acquired at the confocal microscope as described, taking care not to saturate the signal. A maximum intensity projection of 7 slices from each z-stack was thresholded in the *γ*-H2AX channel for both conditions. The threshold was kept the same for all images. Then the Analyze Particles plugin in Fiji was used to quantify all particles of pixel size >30 within this threshold for Mean Intensity and Area. The number of nuclei in each image were counted manually to obtain the number of foci per nucleus.

Measurement of MT growth speed and duration was performed manually in movies of cells expressing EB3-GFP using the MtrackJ plugin in Fiji ^106^. Plus-ends were measured from cells acquired from 3 independent experiments. Measurement of EB1- and CLIP170-labelled plus-ends in fixed cells stained with respective antibodies was performed by taking 2-3 slices from individual z-stacks and generating a maximum intensity projection (MIP). In this MIP, the segmented line ROI (width =3) was used to outline plus-ends. The number of slices in the MIP was kept constant for all images within an experiment. The Measure tool was then used to calculate lengths and mean intensity. To measure the comet density, we outlined an area ROI in a single slice from a z-stack and counted all plus-ends within this ROI. 3-4 ROIs were counted for each image. Measurement of Spindle pole lengths were made using anti-acetylated tubulin stainings in methanol fixed cells. Pole length was manually outlined using the Line ROI tool. For KIF2A stainings, each of the two spindle poles stained with KIF2A was manually outlined using the Segmented Area ROI in Fiji. The average intensity of the two was then taken as the pole intensity for a given cell.

All quantifications were performed on raw, unprocessed images. Maximum intensity projections (MIPs) were generated in Fiji ^107^. Fiji. Images were prepared for publication using Fiji and Adobe Illustrator.

### Statistical analysis

Details of statistical analyses are included in all relevant figure legends. In general, experiments were repeated independently at least twice for biological reproducibility. The number of technical replicates (n) and biological replicates (N) are distinguished in the figure legends for each experiment. For all pairwise comparisons, the unpaired t-test was used. For comparing multiple groups, we used one-way ANOVA followed by the Tukey’s post hoc test. Statistical tests were performed in R. P-values are shown in the figures.

**Supplementary Video 1**

Timelapse video of MT plus-ends labelled with EB3-GFP in 293F cells, stably expressing DOX-inducible dual tubulin, without DOX added (ie. control). Images were acquired on a confocal spinning disk microscope with a frame rate of 0.5 sec, and a pixel resolution of 67 nm. The movie has been cropped and accelerated to 10 frames per second. Image smoothing was performed in Fiji for better visualization.

**Supplementary Video 2**

Timelapse video of MT plus-ends labelled with EB3-GFP in 293F cells, stably expressing DOX-inducible dual tubulin, with 2 *μ*g/ml DOX added for 24 hours prior to imaging (i.e DOX). All other parameters are identical to Supplementary Video 1.

**Supplementary Video 3**

Timelapse video of control cells (without DOX) stained with Sir-Tubulin to visualize mitotic events and imaged on an InviSPIM fluorescent light sheet microscope with a 633 nm laser. Images were acquired every 3 min. Images have been cropped from the original size to zoom in on a single mitotic event, and accelerated to 10 frames per second. Image smoothing was performed in Fiji for better visualization.

**Supplementary Video 4**

Timelapse video of DOX-treated cells stained with Sir-Tubulin to visualize mitotic events and imaged on an InviSPIM fluorescent light sheet microscope with a 633 nm laser. Note the multipolar spindle at the onset of prometaphase. All other parameters are identical to Supplementary Video 3.

**Supplementary Table 1.**

Table showing respectively: normalised counts of the 24 RNA-seq experiments (sheet 2), DESeq2 analysis to determine DEGs after 24 hr of Dox-induction (sheet 3), Metascape analysis of DEGs after 24 hr of Dox-induction (sheet 4), GSEA of the complete RNA-seq dataset (sheet 5), DESeq2 analysis to determine DEGs after 48 hr of Dox-induction (sheet 6), Proteomics analysis of Dox-induced and control cells (sheet 7), K-means clustering analysis of proteomics dataset (sheet 8), Metascape analysis of DEPs after 24 hr of Dox-induction (sheet 9), IPA analysis of DEPs (stringent) (sheet 10), IPA analysis of extended dataset (relaxed) (sheet 11), IPA - specific pathways (sheet 12), TAPs identified after ALFA pull down of HEK293F lysates (sheet 13), TAPs identified after biotin-steptavidin pull down of HeLa cell lysates (sheet 14).

---

**Figure S1.**
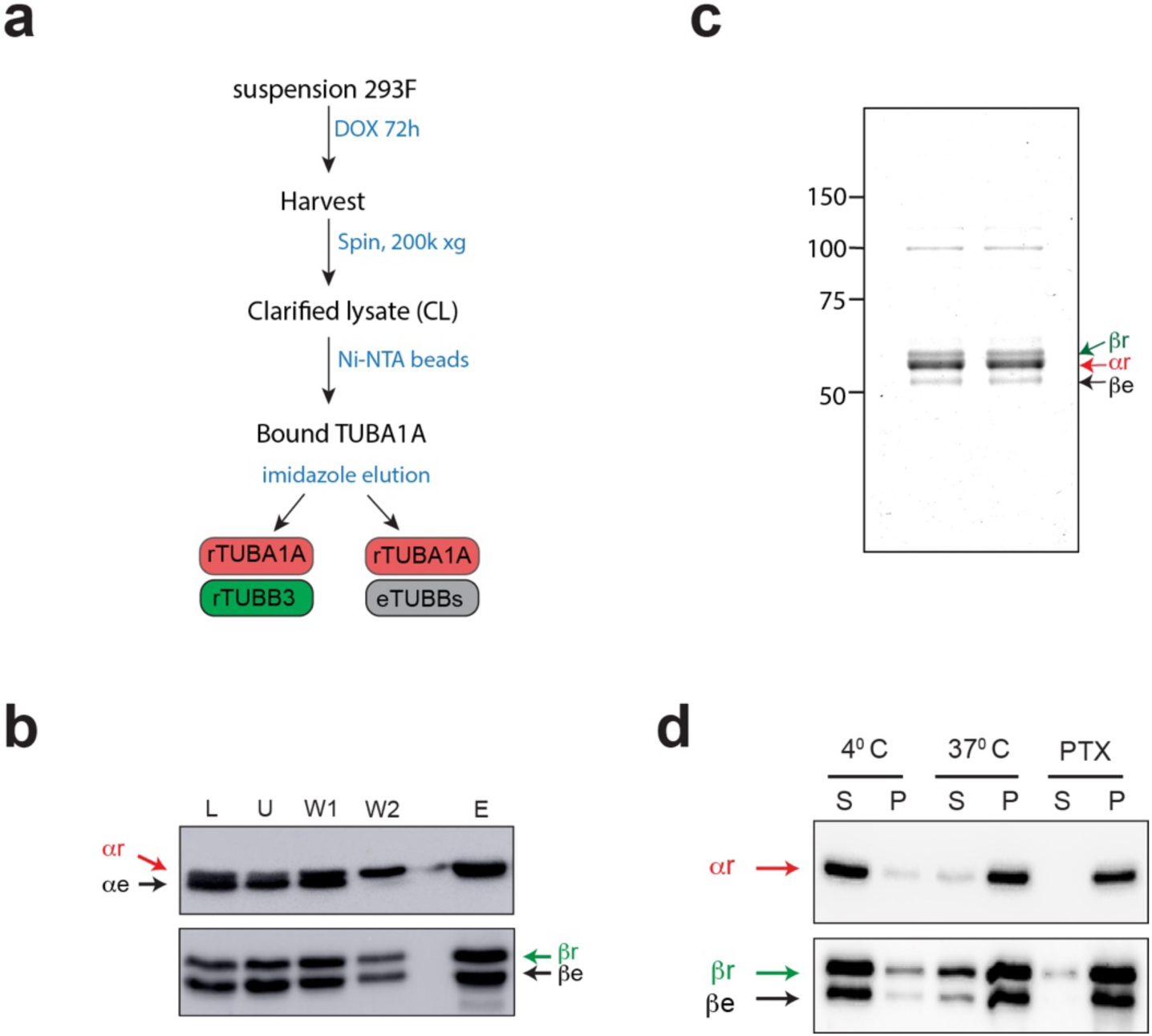
Purification of recombinant tubulin. **a)** Purification strategy of recombinant tubulin from suspension culture after 72 hr of Dox-induction. **b)** Western blots showing the purification of recombinant alpha-tubulin (ar, upper blot) and co-purification of recombinant (br) and endogenous (be) beta-tubulins using the procedure depicted in (A). L: lysate, U: unbound material, W1: first wash, W2: second wash, E: eluate. **c)** Coomassie-stained SDS gel of purified recombinant α/β tubulin, arrowheads indicate the recombinant (r) and endogenous (e) proteins. The position of molecular weight markers is indicated to the left in kilodaltons. **d)** MT pelleting assay under specified conditions showing the fraction of functional recombinant tubulin incorporated into MTs. Note that recombinant α dimerises with both endogenous and recombinant mammalian β tubulin.

**Figure S2.**
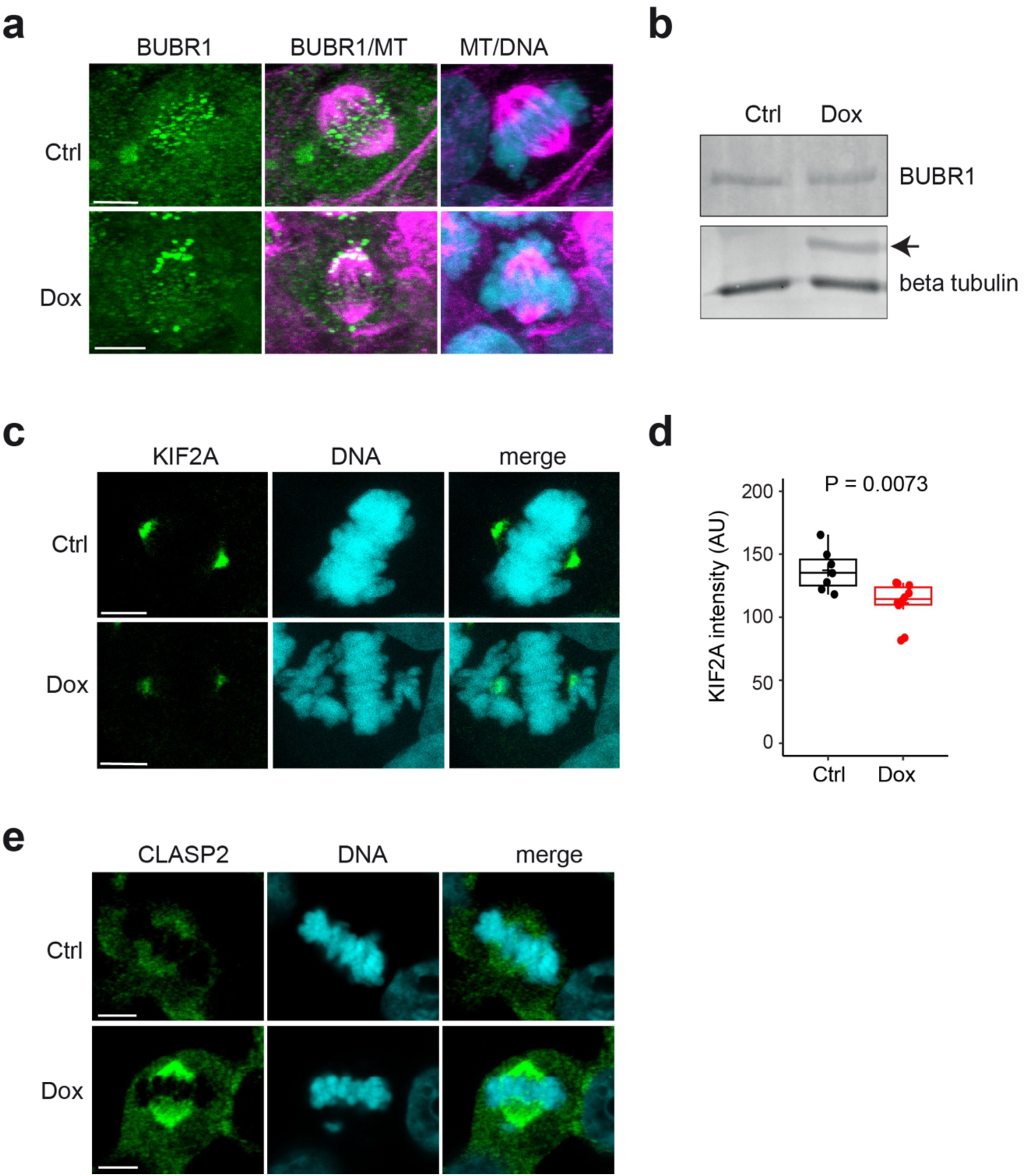
Tubulin overexpression leads to mitotic defects. **a)** BUBR1 localisation in mitotic 293F cells. Cells were treated with Doxycycline (Dox) for 24 hours, or not treated (Ctrl). Cells were fixed, and stained with antibodies against BUBR1 (green) or tubulin (MT, magenta), and with DAPI (cyan) to visualise DNA. Scale bar = 5 *μ*m. **b)** BUBR1 levels in 293F cells. Cells were treated with Doxycycline (Dox) for 24 hours, or not treated (Ctrl). Cells were lysed and total cell lysates analysed by SDS-PAGE and western blot using the indicated antibodies. The arrow indicates the position of recombinant tubulin. A representative western blot is shown (n = 2). **c)** KIF2A localisation in mitotic 293F cells. Cells were treated with Doxycycline (Dox) for 24 hours, or not treated (Ctrl). Cells were fixed, and stained with antibodies against KIF2A (green) and with DAPI (cyan) to visualise DNA. Scale bar = 5 *μ*m. **d)** Quantification of KIF2A intensity. The KIF2A intensity was measured at the spindle pole in cells as shown in (**c**). n= 7 ctrl and 10 Dox spindles. Student T-Tests reveal significant differences between Ctrl and Dox. **e)** CLASP2 localisation in mitotic 293F cells. Cells were treated with Doxycycline (Dox) for 24 hours, or not treated (Ctrl). Cells were fixed, and stained with antibodies against CLASP2 (green) and with DAPI (cyan) to visualise DNA. Scale bar = 5 *μ*m.

**Figure S3.**
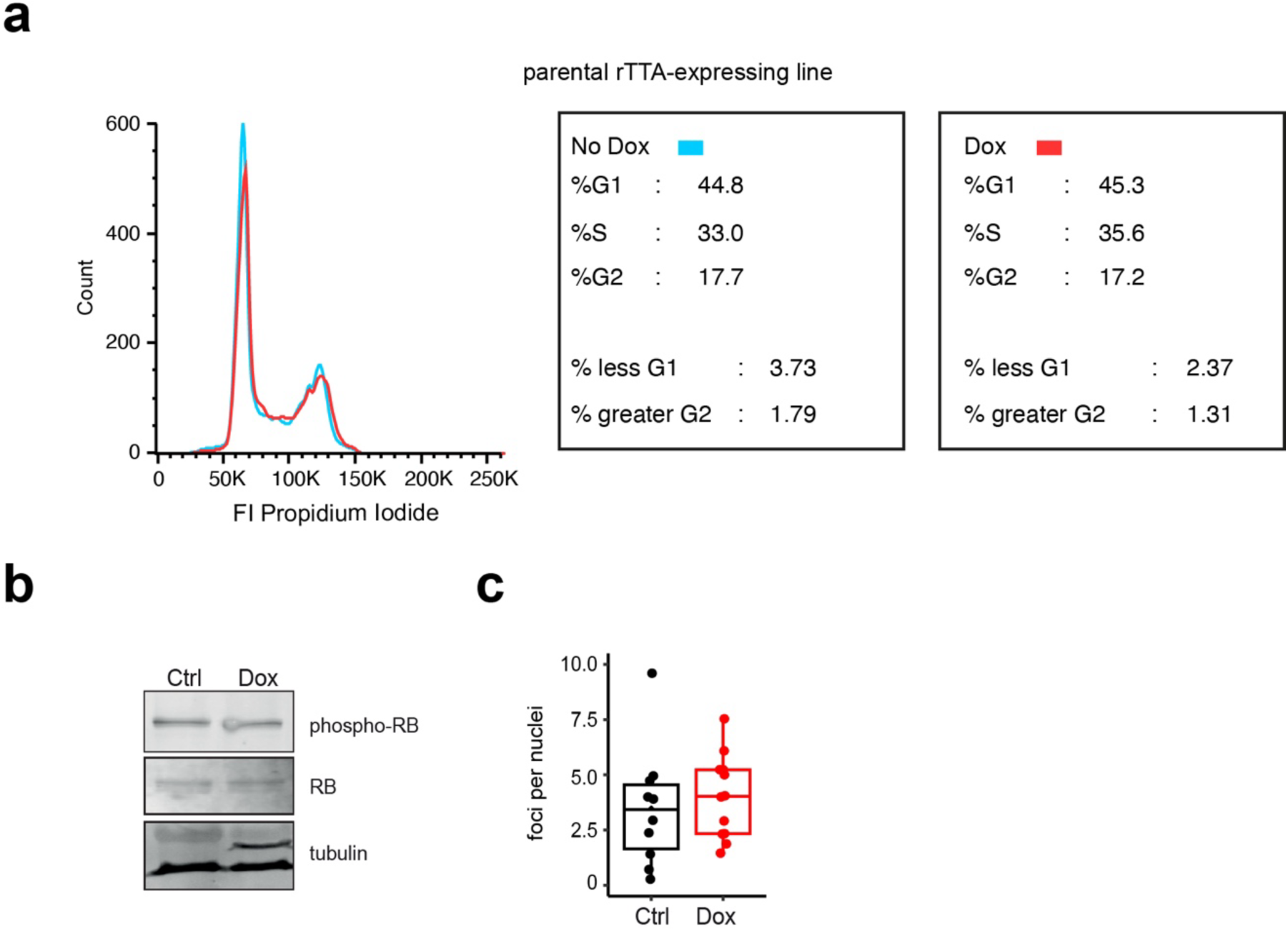
Effects of Doxycycline treatment. **a)** Cell cycle analysis of 293F cells. The parental rTTA-expressing 293F cell line was treated with Doxycycline (Dox, red) for 24 hours, or not treated (No Dox, blue). Cells were fixed, stained with propidium iodide, and analysed by flow cytometry. The left panel shows a representative propidium iodide profile. FI: fluorescence intensity. The middle and right panels show the fractions of cell cycle phases (in percentage). **b)** RB phosphorylation after Dox-induction. Cells were treated with Dox for 24 hours, or not treated (Ctrl). Cells were lysed and total cell lysates analysed by SDS-PAGE and western blot using the indicated antibodies. A representative western blot of RB phosphorylation is shown. **c)** *γ*-H2AX localisation in 293F cells. Cells were treated with Dox for 24 hours, or not treated (Ctrl). Cells were fixed, and stained with anti-*γ*-H2AX antibodies and DAPI to visualise DNA. Maximum intensity projections of 7 slices were analysed (see Fig. 3e for still images). Foci: n =10 control images; 179 nuclei, and 12 Dox-induced images; 246 nuclei, 2 independent experiments. An unpaired T-test revealed no significant differences between Ctrl and Dox.

**Figure S4.**
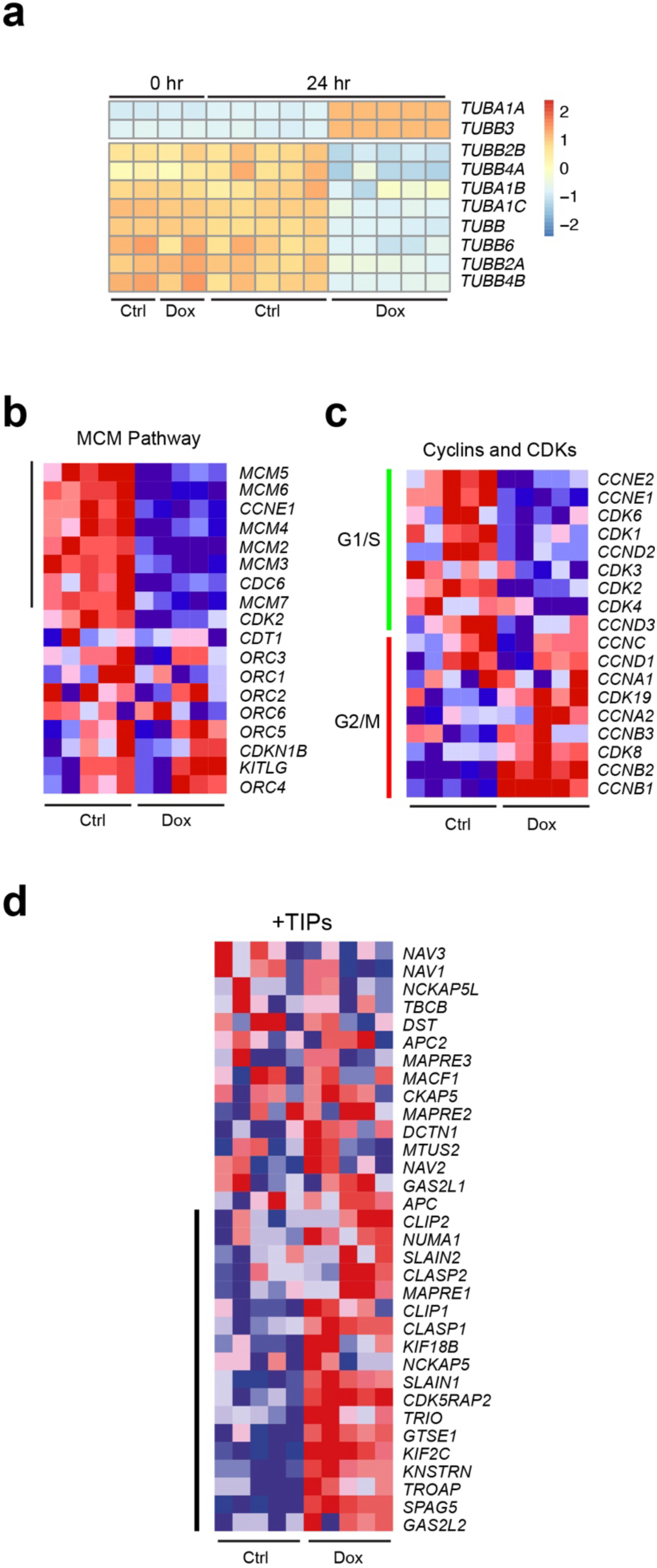
Transcriptomic effect of 24 hr tubulin overexpression. **a)** Heatmap of tubulin expression. 293F cells were treated with Dox for the indicated amount of time, or not treated (Ctrl). mRNA was isolated from the cells and sequenced. The levels of tubulin-encoding mRNAs are shown using a heatmap representation. **b-d**) Heatmaps of gene expression in selected pathways. Gene Set Enrichment Analysis (GSEA) after 24 hr Dox-induction revealed that the MCM pathway involved in DNA replication was significantly down in Dox-induced cells (**b**). In addition, G1/S factors were downregulated in Dox-induced cells in a locally assembled set of mRNAs encoding cyclins and cyclin-dependent kinases (**c**, green line), whereas cyclins and cyclin-dependent kinases important for G2/M were upregulated (**c**, red line). Moreover, and that many +TIPs were upregulated after 24 hr Dox-induction in Dox-induced samples (**d**). Black lines in (**b**) and (**d**) indicate genes that contribute to the enrichment score).

**Figure S5.**
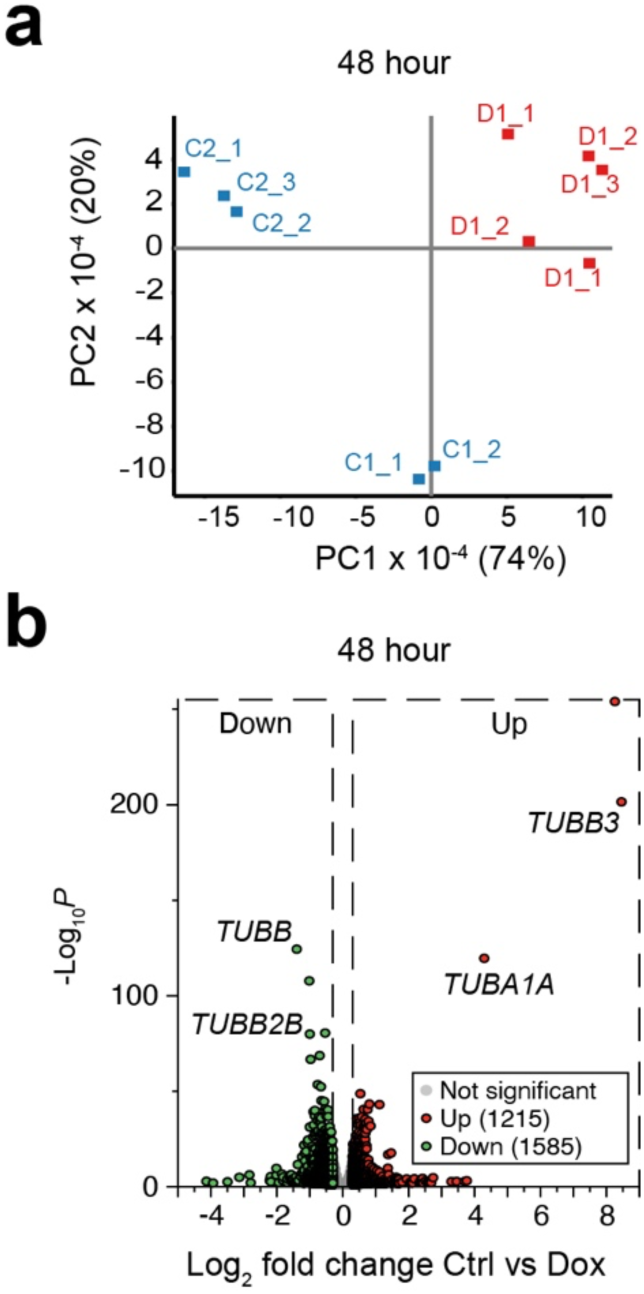
Transcriptomic effect of 48 hr tubulin overexpression. **a)** Principal component analysis (PCA) performed on RNA samples derived from control cells (C, in blue) or Dox-induced cells (D, in red) cultured for 48 hr. Two independent experiments were performed, in the first duplicate samples were analysed (C1_1, C1_2, D1_1, D1_2), whereas in the second triplicates were analysed (C2_1, C2_2, C2_3, D2_1, D2_2, D2_3). **b)** Volcano plot of differentially expressed genes after 48 hr Doxycycline (Dox)-treatment. Cells were either not treated (Ctrl) or treated with Dox for 48 hr. Green and red points indicate differentially expressed mRNAs (green: down in Dox, red: up in Dox) with p-value < 0.05 and a Log2 fold change of less than - 0.3 (green) or more than 0.3 (red). Selected tubulin-encoding mRNAs are indicated.

**Figure S6.**
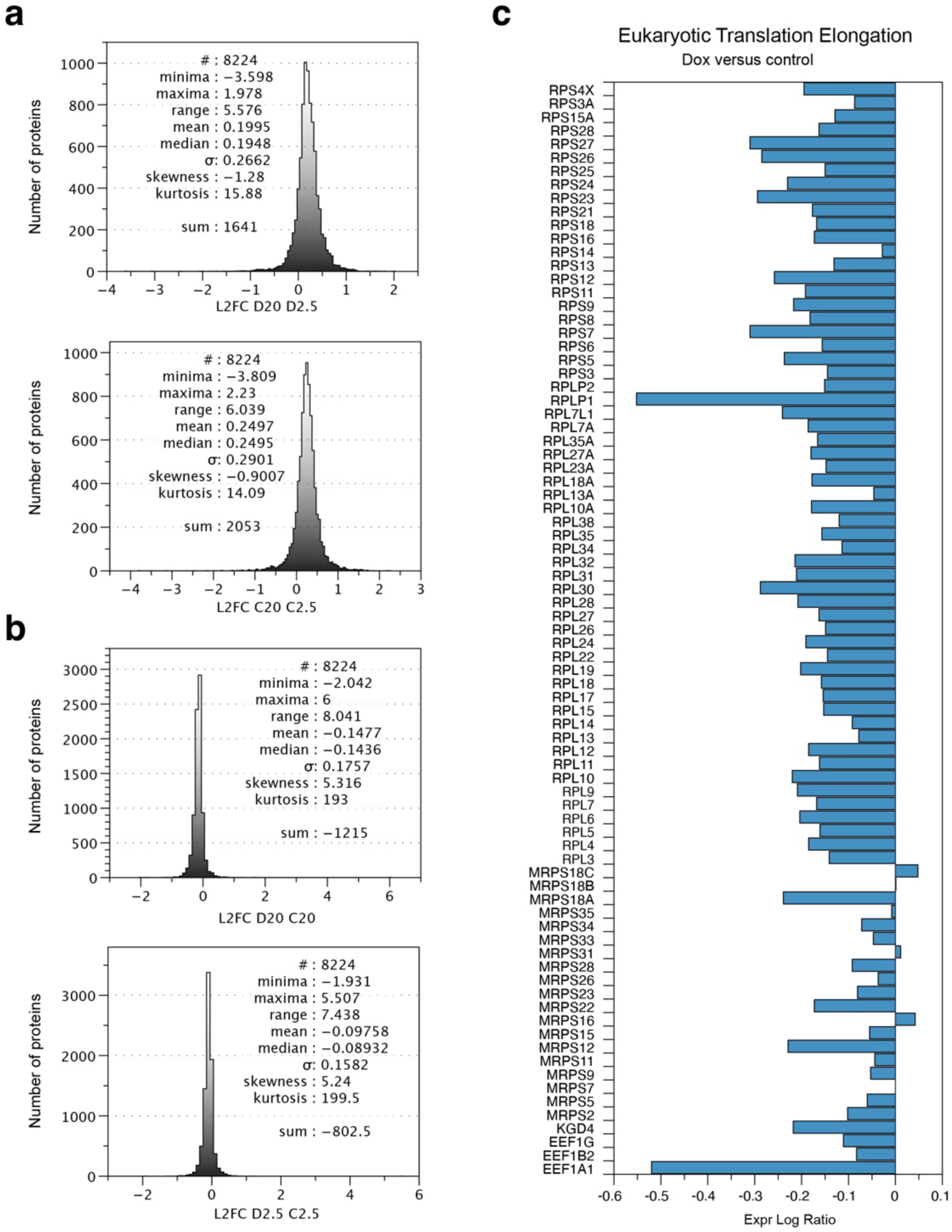
Proteome skewing after hypoxia and tubulin overexpression. **a**, **b**) Probability distributions of proteomes comparing normoxia (20) to hypoxia (2.5) and Dox-induced cells (D) to control cells (C). In (**a**) normoxia is compared to hypoxia (left panel, control cells, right panel, Dox-induced cells), in (**b**) Dox-induced cells are compared to control cells (left panel, hypoxia, right panel, normoxia). In all samples skewness deviates from 0, indicating asymmetry. **c)** Ingenuity Pathway Analysis (IPA) of the Eukaryotic Translation Elongation pathway. Differential expression (in Expr Log Ratio) of proteins of the Eukaryotic Translation Elongation pathway is plotted (see Table S1 for exact values). The more negative the difference, the lower the expression in Dox-induced cells (Dox). Notice how virtually all proteins are expressed at lower level in Dox-induced cells.

**Figure S7.**
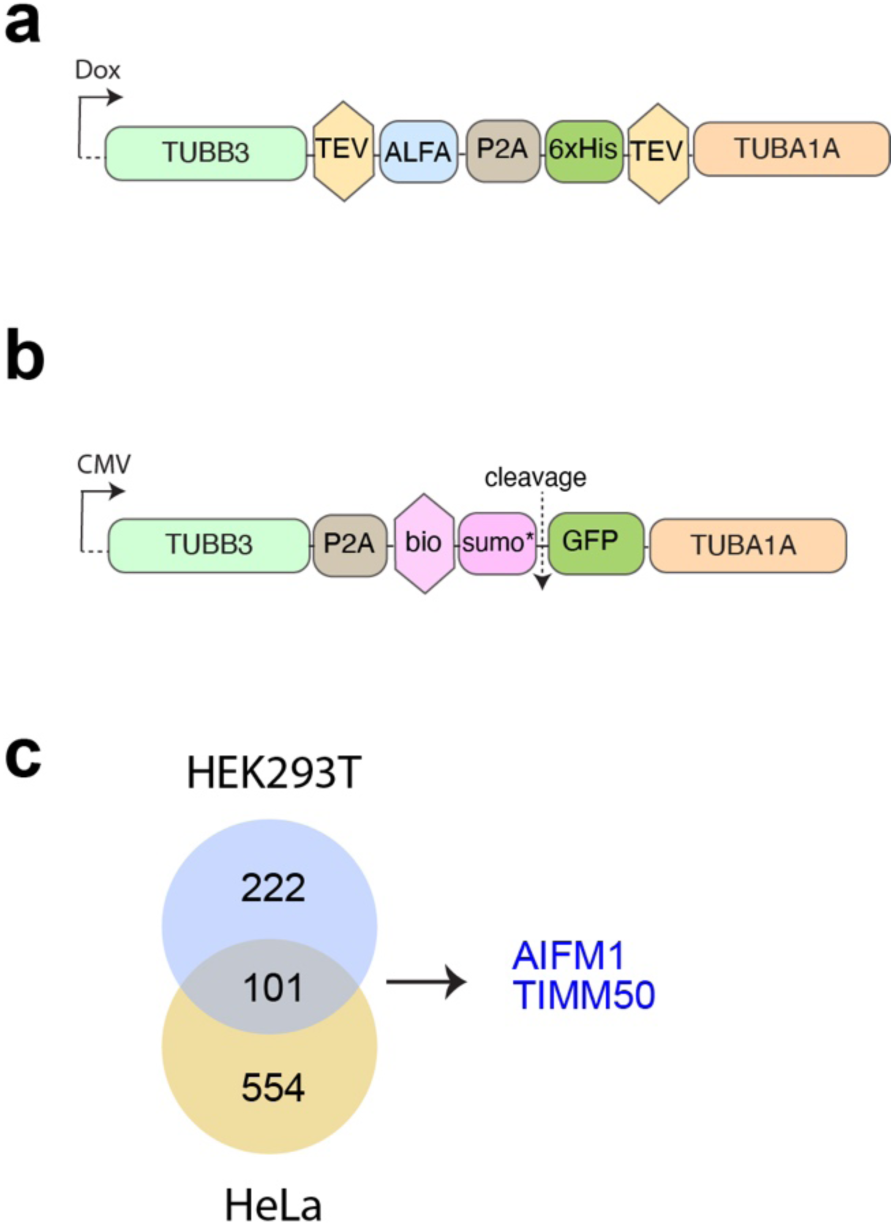
The tubulin interactome of 293F cells. **a**, **b**) Schematic representations of dual tubulin constructs used for tubulin-TAP pull downs. In (**a**) the construct is shown used for pull downs in stably expressing 293F cells (see Figure 7 for results). This dual recombinant tubulin construct is expressed under Tet-responsive promoter (Dox). In (**b**) the construct is shown used for pull downs in transiently transfected HEK293T and HeLa cells (see (**c**) for results). This dual recombinant tubulin construct is expressed under control of the CMV promoter/enhancer (CMV). TEV: TEV protease cleavage site, ALFA: ALFA tag, P2A: the P2A self-cleaving peptide sequence, 6xHis: six consecutive histidines, bio: biotinylation sequence, sumo*: peptide specifically recognised by the Sumo* protease, which cleaves right after this peptide (cleavage). **c)** Analysis of tubulin-associated proteins (TAPs). HEK293T or HeLa cells were transiently transfected with the dual tubulin constructs depicted in (**b**). TAPs were identified using a dual mass spectrometry approach (see main text and Methods for details). AIFM1, and TIMM50 are present in the common pool of TAPs (see Table S1).

## Notes

### Competing Interest Statement

The authors have declared no competing interest.

### Summary of Updates

A revision plan is included based on the comments of the reviewers. No changes were made to the manuscript or the supplemental files.

## References

1. Brouhard, G.J. & Rice, L.M. Microtubule dynamics: an interplay of biochemistry and mechanics. Nat Rev Mol Cell Biol 19, 451–463 (2018).

2. Desai, A. & Mitchison, T.J. Microtubule polymerization dynamics. Annu Rev Cell Dev Biol 13, 83–117 (1997).

3. Goodson, H.V. & Jonasson, E.M. Microtubules and Microtubule-Associated Proteins. Cold Spring Harb Perspect Biol 10 (2018).

4. Breuss, M. & Keays, D.A. Microtubules and neurodevelopmental disease: the movers and the makers. Adv Exp Med Biol 800, 75–96 (2014).

5. Cleveland, D.W. Autoregulated control of tubulin synthesis in animal cells. Curr Opin Cell Biol 1, 10–14 (1989).

6. Steinmetz, M.O. & Prota, A.E. Microtubule-Targeting Agents: Strategies To Hijack the Cytoskeleton. Trends Cell Biol 28, 776–792 (2018).

7. Steinmetz, M.O. & Prota, A.E. Structure-based discovery and rational design of microtubule-targeting agents. Curr Opin Struct Biol 87, 102845 (2024).

8. Ben-Ze’ev, A., Farmer, S.R. & Penman, S. Mechanisms of regulating tubulin synthesis in cultured mammalian cells. Cell 17, 319–325 (1979).

9. Yen, T.J., Machlin, P.S. & Cleveland, D.W. Autoregulated instability of beta-tubulin mRNAs by recognition of the nascent amino terminus of beta-tubulin. Nature 334, 580–585 (1988).

10. Lin, Z. et al. TTC5 mediates autoregulation of tubulin via mRNA degradation. Science 367, 100–104 (2020).

11. Hopfler, M. et al. Mechanism of ribosome-associated mRNA degradation during tubulin autoregulation. Mol Cell 83, 2290–2302 e2213 (2023).

12. Batiuk, A. et al. Soluble alphabeta-tubulins reversibly sequester TTC5 to regulate tubulin mRNA decay. Nat Commun 15, 9963 (2024).

13. Yu, N. et al. Isolation of Functional Tubulin Dimers and of Tubulin-Associated Proteins from Mammalian Cells. Curr Biol 26, 1728–1736 (2016).

14. Yu, N. & Galjart, N. TAPping into the treasures of tubulin using novel protein production methods. Essays Biochem 62, 781–792 (2018).

15. Urlinger, S. et al. Exploring the sequence space for tetracycline-dependent transcriptional activators: novel mutations yield expanded range and sensitivity. Proc Natl Acad Sci U S A 97, 7963–7968 (2000).

16. Aillaud, C. et al. Vasohibins/SVBP are tubulin carboxypeptidases (TCPs) that regulate neuron differentiation. Science 358, 1448–1453 (2017).

17. L’Hernault, S.W. & Rosenbaum, J.L. Chlamydomonas alpha-tubulin is posttranslationally modified in the flagella during flagellar assembly. J Cell Biol 97, 258–263 (1983).

18. Nieuwenhuis, J. et al. Vasohibins encode tubulin detyrosinating activity. Science 358, 1453–1456 (2017).

19. Schroder, H.C., Wehland, J. & Weber, K. Purification of brain tubulin-tyrosine ligase by biochemical and immunological methods. J Cell Biol 100, 276–281 (1985).

20. Hammond, J.W. et al. Posttranslational modifications of tubulin and the polarized transport of kinesin-1 in neurons. Mol Biol Cell 21, 572–583 (2010).

21. Stepanova, T. et al. Visualization of microtubule growth in cultured neurons via the use of EB3-GFP (end-binding protein 3-green fluorescent protein). J Neurosci 23, 2655–2664 (2003).

22. Wood, L.M. & Moore, J.K. beta3 accelerates microtubule plus end maturation through a divergent lateral interface. Mol Biol Cell 36, ar36 (2025).

23. Kok, M., Huber, F., Kalisch, S.M. & Dogterom, M. EB3-informed dynamics of the microtubule stabilizing cap during stalled growth. Biophys J (2024).

24. Cho, N.H. et al. OpenCell: Endogenous tagging for the cartography of human cellular organization. Science 375, eabi6983 (2022).

25. Folker, E.S., Baker, B.M. & Goodson, H.V. Interactions between CLIP-170, tubulin, and microtubules: implications for the mechanism of Clip-170 plus-end tracking behavior. Mol Biol Cell 16, 5373–5384 (2005).

26. Honnappa, S. et al. Key interaction modes of dynamic +TIP networks. Mol Cell 23, 663–671 (2006).

27. Mishima, M. et al. Structural basis for tubulin recognition by cytoplasmic linker protein 170 and its autoinhibition. Proc Natl Acad Sci U S A 104, 10346–10351 (2007).

28. Boyko, S., Li, Q., Surewicz, K. & Surewicz, W.K. Distinct liquid-liquid phase separation properties of end-binding proteins EB1 and EB3. J Biol Chem, 110849 (2025).

29. Meier, S.M. et al. Multivalency ensures persistence of a +TIP body at specialized microtubule ends. Nat Cell Biol 25, 56–67 (2023).

30. Meier, S.M., Steinmetz, M.O. & Barral, Y. Microtubule specialization by +TIP networks: from mechanisms to functional implications. Trends Biochem Sci 49, 318–332 (2024).

31. Borisy, G. et al. Microtubules: 50 years on from the discovery of tubulin. Nat Rev Mol Cell Biol 17, 322–328 (2016).

32. Hosea, R., Hillary, S., Naqvi, S., Wu, S. & Kasim, V. The two sides of chromosomal instability: drivers and brakes in cancer. Signal Transduct Target Ther 9, 75 (2024).

33. Schmidt, M. & Medema, R.H. Exploiting the compromised spindle assembly checkpoint function of tumor cells: dawn on the horizon? Cell Cycle 5, 159–163 (2006).

34. Lukinavicius, G. et al. Fluorogenic probes for live-cell imaging of the cytoskeleton. Nat Methods 11, 731–733 (2014).

35. Tomer, R., Khairy, K. & Keller, P.J. Light sheet microscopy in cell biology. Methods Mol Biol 931, 123–137 (2013).

36. Musacchio, A. & Salmon, E.D. The spindle-assembly checkpoint in space and time. Nat Rev Mol Cell Biol 8, 379–393 (2007).

37. Jordan, M.A., Thrower, D. & Wilson, L. Effects of vinblastine, podophyllotoxin and nocodazole on mitotic spindles. Implications for the role of microtubule dynamics in mitosis. J Cell Sci 102 **(Pt** **3**), 401–416 (1992).

38. Jordan, M.A., Toso, R.J., Thrower, D. & Wilson, L. Mechanism of mitotic block and inhibition of cell proliferation by taxol at low concentrations. Proc Natl Acad Sci U S A 90, 9552–9556 (1993).

39. Sudakin, V., Chan, G.K. & Yen, T.J. Checkpoint inhibition of the APC/C in HeLa cells is mediated by a complex of BUBR1, BUB3, CDC20, and MAD2. J Cell Biol 154, 925–936 (2001).

40. Logarinho, E. & Bousbaa, H. Kinetochore-microtubule interactions “in check” by Bub1, Bub3 and BubR1: The dual task of attaching and signalling. Cell Cycle 7, 1763–1768 (2008).

41. Gaetz, J. & Kapoor, T.M. Dynein/dynactin regulate metaphase spindle length by targeting depolymerizing activities to spindle poles. J Cell Biol 166, 465–471 (2004).

42. Ganem, N.J. & Compton, D.A. The KinI kinesin Kif2a is required for bipolar spindle assembly through a functional relationship with MCAK. J Cell Biol 166, 473–478 (2004).

43. Uehara, R. et al. Aurora B and Kif2A control microtubule length for assembly of a functional central spindle during anaphase. J Cell Biol 202, 623–636 (2013).

44. Girao, H. et al. CLASP2 binding to curved microtubule tips promotes flux and stabilizes kinetochore attachments. J Cell Biol 219 (2020).

45. Logarinho, E. et al. CLASPs prevent irreversible multipolarity by ensuring spindle-pole resistance to traction forces during chromosome alignment. Nat Cell Biol 14, 295–303 (2012).

46. Maffini, S. et al. Motor-independent targeting of CLASPs to kinetochores by CENP-E promotes microtubule turnover and poleward flux. Curr Biol 19, 1566–1572 (2009).

47. Zhai, Y., Kronebusch, P.J., Simon, P.M. & Borisy, G.G. Microtubule dynamics at the G2/M transition: abrupt breakdown of cytoplasmic microtubules at nuclear envelope breakdown and implications for spindle morphogenesis. J Cell Biol 135, 201–214 (1996).

48. Zhai, Y. & Borisy, G.G. Quantitative determination of the proportion of microtubule polymer present during the mitosis-interphase transition. J Cell Sci 107 **(Pt** **4**), 881–890 (1994).

49. Travis, S.M., Mahon, B.P. & Petry, S. How Microtubules Build the Spindle Branch by Branch. Annu Rev Cell Dev Biol 38, 1–23 (2022).

50. Zhou, A.S. et al. Diverse microtubule-targeted anticancer agents kill cells by inducing chromosome missegregation on multipolar spindles. PLoS Biol 21, e3002339 (2023).

51. Helin, K. Regulation of cell proliferation by the E2F transcription factors. Curr Opin Genet Dev 8, 28–35 (1998).

52. Henley, S.A. & Dick, F.A. The retinoblastoma family of proteins and their regulatory functions in the mammalian cell division cycle. Cell Div 7, 10 (2012).

53. Weinberg, R.A. The retinoblastoma protein and cell cycle control. Cell 81, 323–330 (1995).

54. Rogakou, E.P., Pilch, D.R., Orr, A.H., Ivanova, V.S. & Bonner, W.M. DNA double-stranded breaks induce histone H2AX phosphorylation on serine 139. J Biol Chem 273, 5858–5868 (1998).

55. Sedelnikova, O.A., Pilch, D.R., Redon, C. & Bonner, W.M. Histone H2AX in DNA damage and repair. Cancer Biol Ther 2, 233–235 (2003).

56. Mazumdar, M., Sung, M.H. & Misteli, T. Chromatin maintenance by a molecular motor protein. Nucleus 2, 591–600 (2011).

57. Dullovi, A. et al. Microtubule-associated proteins MAP7 and MAP7D1 promote DNA double-strand break repair in the G1 cell cycle phase. iScience 26, 106107 (2023).

58. Luessing, J. et al. The nuclear kinesin KIF18B promotes 53BP1-mediated DNA double-strand break repair. Cell Rep 35, 109306 (2021).

59. Zhu, S. et al. Kinesin Kif2C in regulation of DNA double strand break dynamics and repair. Elife 9 (2020).

60. Ma, S. et al. DNA damage promotes microtubule dynamics through a DNA-PK-AKT axis for enhanced repair. J Cell Biol 220 (2021).

61. Liu, P., Wurtz, M., Zupa, E., Pfeffer, S. & Schiebel, E. Microtubule nucleation: The waltz between gamma-tubulin ring complex and associated proteins. Curr Opin Cell Biol 68, 124–131 (2021).

62. Gasic, I., Boswell, S.A. & Mitchison, T.J. Tubulin mRNA stability is sensitive to change in microtubule dynamics caused by multiple physiological and toxic cues. PLoS Biol 17, e3000225 (2019).

63. Assaf, M., Lacheheub, C. & Gasic, I. Tubulin autoregulation controls the biosynthesis of γ-tubulin to ensure mitotic fidelity. bioRxiv (2026).

64. Zhou, Y. et al. Metascape provides a biologist-oriented resource for the analysis of systems-level datasets. Nat Commun 10, 1523 (2019).

65. Liberzon, A. et al. The Molecular Signatures Database (MSigDB) hallmark gene set collection. Cell Syst 1, 417–425 (2015).

66. Subramanian, A. et al. Gene set enrichment analysis: a knowledge-based approach for interpreting genome-wide expression profiles. Proc Natl Acad Sci U S A 102, 15545–15550 (2005).

67. Jordan, M.A. & Kamath, K. How do microtubule-targeted drugs work? An overview. Curr Cancer Drug Targets 7, 730–742 (2007).

68. Semenza, G.L., Shimoda, L.A. & Prabhakar, N.R. Regulation of gene expression by HIF-1. Novartis Found Symp 272, 2–8; discussion 8-14, 33-16 (2006).

69. Yfantis, A. et al. Transcriptional Response to Hypoxia: The Role of HIF-1-Associated Co-Regulators. Cells 12 (2023).

70. Costa-Mattioli, M. & Walter, P. The integrated stress response: From mechanism to disease. Science 368 (2020).

71. Chun, Y. & Kim, J. AMPK-mTOR Signaling and Cellular Adaptations in Hypoxia. Int J Mol Sci 22 (2021).

72. Harding, H.P. et al. An integrated stress response regulates amino acid metabolism and resistance to oxidative stress. Mol Cell 11, 619–633 (2003).

73. Amiri, M., Toboz, P., Bellucci, M.A., Tahmasebi, S. & Sonenberg, N. From the integrated stress response to oxidative stress: A historical perspective. J Biol Chem 302, 110958 (2026).

74. Place, T.L., Domann, F.E. & Case, A.J. Limitations of oxygen delivery to cells in culture: An underappreciated problem in basic and translational research. Free Radic Biol Med 113, 311–322 (2017).

75. Kramer, A., Green, J., Pollard, J., Jr. & Tugendreich, S. Causal analysis approaches in Ingenuity Pathway Analysis. Bioinformatics 30, 523–530 (2014).

76. Stepkowski, T.M. et al. Temporal alterations of the nascent proteome in response to mitochondrial stress. Cell Rep 43, 114803 (2024).

77. Gotzke, H. et al. The ALFA-tag is a highly versatile tool for nanobody-based bioscience applications. Nat Commun 10, 4403 (2019).

78. Hangen, E. et al. Interaction between AIF and CHCHD4 Regulates Respiratory Chain Biogenesis. Mol Cell 58, 1001–1014 (2015).

79. Chaudhuri, M., Tripathi, A. & Gonzalez, F.S. Diverse Functions of Tim50, a Component of the Mitochondrial Inner Membrane Protein Translocase. Int J Mol Sci 22 (2021).

80. Carbonaro, M., Escuin, D., O’Brate, A., Thadani-Mulero, M. & Giannakakou, P. Microtubules regulate hypoxia-inducible factor-1alpha protein trafficking and activity: implications for taxane therapy. J Biol Chem 287, 11859–11869 (2012).

81. Akhmanova, A. et al. Clasps are CLIP-115 and -170 associating proteins involved in the regional regulation of microtubule dynamics in motile fibroblasts. Cell 104, 923–935 (2001).

82. Drabek, K. et al. Role of CLASP2 in microtubule stabilization and the regulation of persistent motility. Curr Biol 16, 2259–2264 (2006).

83. Mitchison, T.J. Dynamic microtubules as sensors in animal cells. Biophys J (2025).

84. Vanni, G. et al. Microtubule architecture connects AMOT stability to YAP/TAZ mechanotransduction and Hippo signalling. Nat Cell Biol 27, 1725–1738 (2025).

85. Cleveland, D.W., Lopata, M.A., Sherline, P. & Kirschner, M.W. Unpolymerized tubulin modulates the level of tubulin mRNAs. Cell 25, 537–546 (1981).

86. Mitchison, T.J. The proliferation rate paradox in antimitotic chemotherapy. Mol Biol Cell 23, 1–6 (2012).

87. Mitchison, T.J., Pineda, J., Shi, J. & Florian, S. Is inflammatory micronucleation the key to a successful anti-mitotic cancer drug? Open Biol 7 (2017).

88. Weaver, B.A. How Taxol/paclitaxel kills cancer cells. Mol Biol Cell 25, 2677–2681 (2014).

89. Chuong, S.D. et al. Large-scale identification of tubulin-binding proteins provides insight on subcellular trafficking, metabolic channeling, and signaling in plant cells. Mol Cell Proteomics 3, 970–983 (2004).

90. Gay, D.A., Sisodia, S.S. & Cleveland, D.W. Autoregulatory control of beta-tubulin mRNA stability is linked to translation elongation. Proc Natl Acad Sci U S A 86, 5763–5767 (1989).

91. Pachter, J.S., Yen, T.J. & Cleveland, D.W. Autoregulation of tubulin expression is achieved through specific degradation of polysomal tubulin mRNAs. Cell 51, 283–292 (1987).

92. Komarova, Y. et al. Mammalian end binding proteins control persistent microtubule growth. J Cell Biol 184, 691–706 (2009).

93. Rashid, A., Naaz, A., Rai, A., Chatterji, B.P. & Panda, D. Inhibition of polo-like kinase 1 suppresses microtubule dynamics in MCF-7 cells. Mol Cell Biochem 465, 27–36 (2020).

94. Souphron, J. et al. Purification of tubulin with controlled post-translational modifications by polymerization-depolymerization cycles. Nat Protoc 14, 1634–1660 (2019).

95. Poruchynsky, M.S. et al. Tumor cells resistant to a microtubule-depolymerizing hemiasterlin analogue, HTI-286, have mutations in alpha- or beta-tubulin and increased microtubule stability. Biochemistry 43, 13944–13954 (2004).

96. Kim, D., Paggi, J.M., Park, C., Bennett, C. & Salzberg, S.L. Graph-based genome alignment and genotyping with HISAT2 and HISAT-genotype. Nat Biotechnol 37, 907–915 (2019).

97. Li, H. et al. The Sequence Alignment/Map format and SAMtools. Bioinformatics 25, 2078–2079 (2009).

98. Anders, S., Pyl, P.T. & Huber, W. HTSeq--a Python framework to work with high-throughput sequencing data. Bioinformatics 31, 166–169 (2015).

99. Love, M.I., Huber, W. & Anders, S. Moderated estimation of fold change and dispersion for RNA-seq data with DESeq2. Genome Biol 15, 550 (2014).

100. Mootha, V.K. et al. PGC-1alpha-responsive genes involved in oxidative phosphorylation are coordinately downregulated in human diabetes. Nat Genet 34, 267–273 (2003).

101. Ge, S.X., Son, E.W. & Yao, R. iDEP: an integrated web application for differential expression and pathway analysis of RNA-Seq data. BMC Bioinformatics 19, 534 (2018).

102. Gu, Z. Complex heatmap visualization. Imeta 1, e43 (2022).

103. Gu, Z., Eils, R. & Schlesner, M. Complex heatmaps reveal patterns and correlations in multidimensional genomic data. Bioinformatics 32, 2847–2849 (2016).

104. Schwertman, P. et al. UV-sensitive syndrome protein UVSSA recruits USP7 to regulate transcription-coupled repair. Nat Genet 44, 598–602 (2012).

105. Cox, J. et al. A practical guide to the MaxQuant computational platform for SILAC-based quantitative proteomics. Nat Protoc 4, 698–705 (2009).

106. Meijering, E., Dzyubachyk, O. & Smal, I. Methods for cell and particle tracking. Methods Enzymol 504, 183–200 (2012).

107. Schindelin, J., et al. Fiji: an open-source platform for biological-image analysis. Nat Methods 9, 676–682 (2012).

