## Supplementary material for "A role for tubulin in cellular quality control and proteostasis": Revision plan

**Manuscript number:** RC-2026-03552

**Corresponding author(s):** Niels Galjart

### 1. General Statements

We thank the reviewers for their constructive criticisms and helpful suggestions.

### 2. Description of the planned revisions

*Point-by-point reply explaining what revisions, additional experimentations and analyses are planned to address the points raised by the referees.*

**Reviewer #1**

(Note: reviewer comments are in blue, rebuttal is in black)

Summary: This study examines how simultaneous expression of high levels of alpha and beta tubulin impact microtubule organization, cell cycle progression and stress response. The main conclusions are that a/b tubulin overexpression 1) increases the density of microtubules in cells with mild effects on growth rate and catastrophe frequency; 2) causes mitotic spindle defects and cell cycle disruption; 3) alters the transcriptome and proteostasis; 4) causes mitochondrial stress by binding mitochondrial import proteins; 5) disrupts cellular stress response.

Major comments:

The results generally use an appropriate number of cells and technical replicates.

There are several instances in the manuscript where the data do not support the authors' conclusions. These include:

Page 6. "We detected a mild increase in MT growth rates in Dox-treated cells (Fig. 2b), which was observed in all of the growing MTs examined (Fig. 2c), consistent with an increase in soluble tubulin levels in all Dox-induced cells." The data do not support this statement. The histogram in Figure 2C shows that only a small portion of growth rate measurements are faster in the overexpression cells, and the statistical test in 2b suggests that the datasets are unlikely to be different. It is unclear from the presentation of the data why a small population of overexpression cells exhibit faster growth rates. The data in Figure 2b should be plotted as a superplot could show whether the faster data points are from specific cells or technical replicates. The same concern applies to the data in Figure 2D.

Answer:

We agree that an overlaid density plot as currently shown in Fig. 2c might visually suggest that only the non-overlapping tail differs between conditions, but this is not what the underlying data show. Comparison of individual comet measurements indicates that the entire Dox distribution is shifted toward faster speeds (as we state in the text), not a small subpopulation: a randomly selected Dox comet is faster than a randomly selected Ctrl comet in 62% of pairwise comparisons (50% expected if the groups were identical), and 69% of Dox comets exceed the Ctrl median speed. We have added a cumulative distribution plot (see Figure below) that makes this more explicit — the Dox curve runs to the right of Ctrl across essentially the entire range, rather than only in a discrete high-speed segment. We acknowledge that our previous Figure panel may not have represented the data properly and propose to replace current Fig. 2c with this new plot.

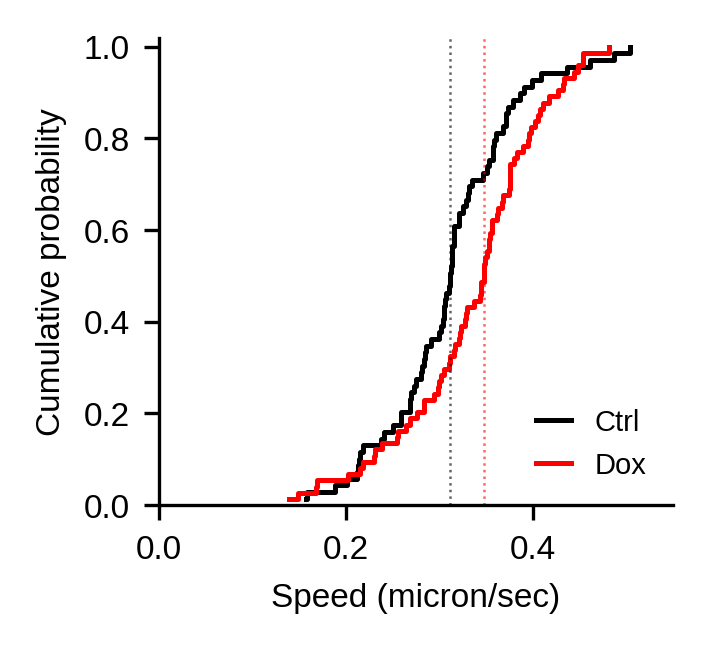

*New Fig. 2c. Cumulative distribution plot of EB3-GFP comets (displacements in time).*

Regarding the statistic in Fig. 2b: the reported P = 0.057 is from a test comparing group means. However, the Dox distribution is right-skewed and borderline non-normal (Shapiro-Wilk P = 0.054), and we realized that this is a condition under which a mean-comparison test is not well powered. A Mann-Whitney U test, which compares the full rank distribution and is more appropriate for data of this shape, gives P = 0.013 (Kolmogorov-Smirnov P = 0.0043). We will add these results and the effect-size measures mentioned above to a revised manuscript, and will report the rank-based test as the primary statistic for this comparison in the revised manuscript. All in all, we hope this clarifies that the effect is a genuine population-wide shift in comet speed. Note that we agree with the reviewer that the data should be plotted as a superplot in Fig. 2b, and we will do this in a revised manuscript.

For the data presented in Fig. 2d (duration of EB3-GFP displacements), we performed a similar analysis as described above. In contrast to Fig. 2c no significant changes are seen in duration, and here the concern of the reviewer is justified. We will indicate this clearly in the text. It does not change any of our conclusions.

Page 11. "Moreover, DNA content was highly aberrant after 48 hr, with large proportions of cells containing < 2n or > 4n chromosomes (Fig. 4d). Thus, persistent overexpression of tubulin affects the cell cycle and after 48 hr it results in severe DNA abnormalities, suggestive of CIN". The plot in Figure 4D shows increased propidium iodide signal below the 2N peak and above the 4N peak, but it is quite likely that this could represent signal from apoptotic cells. The authors should specifically stain for an apoptotic marker to test this possibility. This would suggest that prolonged overexpression of a/b tubulin leads to apoptosis, which could be important evidence for later conclusions in the study.

Answer:

We agree with the reviewer. We actually thought along the same lines and already performed the apoptosis experiment by staining tubulin overexpressing cells (dox) and control cells (ctrl) with Annexin, an apoptosis marker, and propidium iodide. We then quantified double-stained cells. The experiment was performed in triplicate. As shown in the Figure below (averages ± SEM), the data do not reveal differences in apoptosis. We therefore did not include these results in the original manuscript. However, based on the comment of the reviewer we will now include these data in a revised version of the manuscript. The conclusion is that tubulin overexpression does not lead to increased apoptosis.

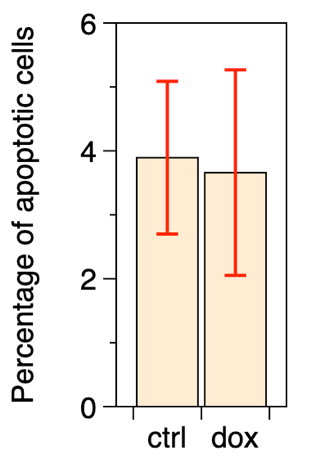

*Apoptosis Figure.*

Page 11. "Combined, our data show that tubulin overexpression affects all three major cell cycle checkpoints, i.e. G1/S, G2/M, and the SAC in mitosis." The data do not fully support this conclusion. While the evidence for SAC-mediated delay in overexpression cells is strong, the evidence for other checkpoints is weak. The phosphor-RB experiment in Figure S3b lacks a positive control. The γH2A-X results in Figure 4f and g show that the difference is driven by a minor population of dim foci in control cells. Similar to the point above, these data should be plotted as a superplot to explore the possibility that this minor population arises from a small number of cells or a specific technical replicate.

Answer:

With respect to the “inclusion of a positive control”, we are not sure what is meant by the reviewer. Total RB is shown underneath the phosphor-RB lane, and underneath that lane we show a tubulin blot. Moreover, mass spectrometry data are included in Table 1, showing the levels of many other proteins. Note that both the western blot and proteomics data reveal similar RB1 levels in Ctrl and Dox cells.

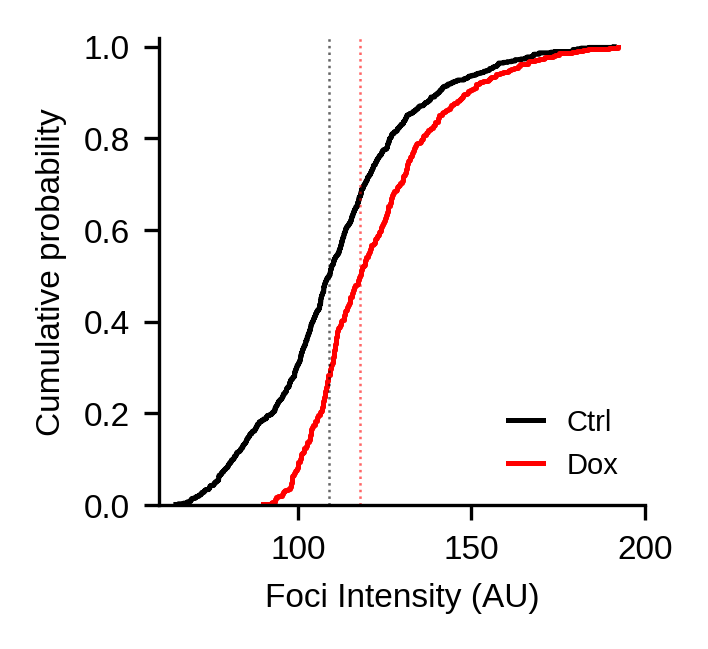

*New Fig. 4g. Cumulative distribution plot of 𝛾-H2AX foci intensity.*

With respect to the γH2A-X results in Figure 4f and g, we re-analyzed and re-plotted the data as described above for Fig. 2c. A cumulative distribution plot, depicted above, clearly shows separation of the two populations. We ran several statistical tests, which all show highly siginificant values (Welch P = 1.5e-17, Mann-Whitney P = 3.2e-17, KS P = 1.1e-14). We propose to change the current Fig. 4g with the plot shown above, and we will indicate statistical tests.

We then tested the reviewer's proposal (*driven by a minor population of dim foci in control cells*) by fitting a 2-component Gaussian mixture to each condition separately. The table below shows close to the opposite of the reviewer's hypothesis. Both Ctrl and Dox contain the same two-population structure in similar proportions (roughly 75%/25%). The minor subpopulation is therefore not unique to Ctrl, but a feature of both conditions. Perhaps it is a cell-cycle-linked fraction. When we next compared the matched components directly we observed that the majority/lower-intensity component is shifted (102 → 112 AU, P = 1.5e-30), while the minor/higher-intensity component is statistically indistinguishable between conditions (143 vs. 146 AU, P = 0.11, ns). Thus, if anything, the effect is carried by the bulk population, and the minor subpopulation is the part that does not differ. All data are summarized in the table below.

| **2-component Gaussian mixture decomposition** | **Ctrl weight** | **Ctrl mean (AU)** | **Dox weight** | **Dox mean (AU)** |
| --- | --- | --- | --- | --- |
| Lower-intensity component | ~78% | 101.7 | ~71% | 111.9 |
| Higher-intensity component | ~22% | 143.3 | ~29% | 146.3 |

Page 19. "This, in turn, lowers EEF1A1 causing attenuation of translation and dampened elongation rates. That other initiation and elongation factors are mildly down in Dox-induced cells, as are ribosomal proteins (Fig S6c, Table S1), indicates a mild ISR and general translation inhibition. We propose that tubulin overexpression results in mitochondrial dysfunction leading to stress and attenuated translation elongation. We observe the start of an ISR in these cells, which prevents a proper hypoxic response." This conclusion seems to be at odds with the earlier conclusion drawn from Figure 5e, where ISR genes are significantly decreased in overexpression cells. The authors should reconcile these results in the discussion.

Answer:

Our proteomics data suggest that attenuated translation elongation caused by lower EEF1A1 levels hampers the integrated stress response (ISR) in Dox-induced cells. One can intuitively understand this: translation is already lowered, it does not need to be lowered much further via the ISR. Thus, in Dox-induced cells the ISR is hampered compared to control cells, which do not suffer from mito-stress. In other words, the ISR is set in motion in Dox-induced cells but less compared to control cells. This is exactly what we see in the RNA-Sequencing experiments, where in Dox-induced cells mRNAs encoding proteins in volved in HIF-1 and PKR signaling (reflecting ISR) are down (Fig. 5e, right hand panel). Thus, ISR is hampered at the RNA level when tubulin is overexpressed. While we thought we had explained our view well enough on page 19, we realize that this may not have been the case and we will provide an improved explanation in the Discussion section of a revised manuscript.

Figure 7. This experiment lacks a key control - identifying peptides in a pull down from cells that do not overexpress the tagged tubulin. Without this control, it is impossible to discern which peptides bind to the beads, independent of tubulin. This concern is amplified by the results in Figure 7b where TIMM50 binding is tested in a follow-up experiment. Here the negative control is employed and it shows that TIMM50 pulls down in the absence of tagged tubulin. This indicates that TIMM50 may not bind to tubulin and calls into question any conclusions related to TIMM50 and mitochondrial proteostasis. Before making any conclusions from the pull down experiment, the authors must include the negative control and carefully assess which peptides are likely to be false positives.

Answer:

In contrast to what the reviewer states a negative control was included. As described on page 19: “Using beads coupled to ALFA-tag antibodies, we affinity-purified recombinant tubulin and tubulin-associated proteins (TAPs) from cell lysates of tubulin overexpressing cells, using non-induced cells as controls”. To make this more clear we propose to include a plot in Figure 7 showing the fold enrichment of TAPs in the dox-induced versus Ctrl cells.

The reason that TIMM50 enrichment in the recombinant tubulin pull down is not high is explained on page 21 of the manuscript: “We note that the relatively weak enrichment of TIMM50 on beads containing recombinant tubulins (Fig. 7b, lower blot, Table S1) as compared to the enrichment of the recombinant tubulins themselves (Fig. 7b, upper and middle blots, Table S1) is well explained, first by a low affinity of the tubulin-TIMM50 interaction itself, and second by competition for TIMM50 between recombinant tubulins, which are enriched after washing, and the much larger reservoir of endogenous tubulins, which are lost (together with TIMM50) after washing”.

While we believe that the weak enrichment of TIMM50 is well explained in the current set-up, we nevertheless feel that the tubulin-TIMM50 interaction should be further corroborated and we propose to perform a new experiment where we tag both TIMM50 (and AIFM) in addition to our dual tubulin constructs and show - via dual affinity purification - that the proteins do interact.

Figure 8. This figure and the associated text feel rather disconnected from the rest of the study. The authors do not make clear conclusions from these results. Perhaps these experiments should be further developed in a separate study?

Answer:

The physiological relevance of autoregulation is still largely unknown. Our analysis in Figure 8, where we show that autoregulation is activated upon stress (i.e. hypoxia and Gln deprivation), provides a clue. Moreover, we show that disruption of normal tubulin levels, which induces stress, dampens the hypoxic response. We believe these reciprocal effects fit nicely. However, since reviewer 2 is of the same opinion as this reviewer we are willing to remove the data in a revised version of the manuscript.

Page 25. "We found that tubulin overexpression induces mitochondrial dysfunction." The data do not strongly support this conclusion. The proteomic data in Figure 6 indicate that some mitochondrial proteins are less abundant in tubulin overexpressing cells, but the study does not include any experiments that actually test mitochondrial function. The authors could test this by including new experiments to measure mitochondrial membrane potential, or respiration activity, etc. These would be an important addition to the study.

Answer:

We agree with this criticism and will perform additional experiments in a revised version of the manuscript to describe mitochondrial dysfunction.

Page 26. "We provide evidence here that surplus tubulin slows general translation, increasing the time window for a TTC5-nascent tubulin interaction." The data do not demonstrate a decrease in translation rate. It is unclear what evidence the authors refer to with this statement. The proteomic analysis in Figure 6 measures protein abundance, and a decrease could be due to either decreased transcription, decreased translation, or increased protein degradation. The number of proteins that are shown to be downregulated in tubulin overexpressing cells is rather small (100s) which seems to argue against a general decrease in translation, despite the decrease in several translation regulators. To make this conclusion, the authors would need to add new experiments that measure translation rate.

Answer:

While we agree with the reviewer that “protein decrease could be due to either decreased transcription, decreased translation, or increased protein degradation”, we provide evidence below that, at least in our view, excludes transcription and degradation as mechanisms. We do agree that we should provide more compelling evidence that attenuated translation is at work in tubulin overexpressing cells. We will therefore perform an assay to measure translation rates in tubulin overexpressing and control cells.

Having stated which new experiment we will perform in a revised version of the manuscript, we now would like to extensively react to the above comment of the reviewer using our existing data. First, the reviewer argues that “the number of proteins that are shown to be downregulated in tubulin overexpressing cells is rather small (100s)”. However, this applies to the signficantly differentially expressed proteins (DEPs) and "only hundreds of proteins pass significance" is a power argument, not an effect-size argument. In Fig. 7c (heatmap) and Fig. S6b (probability distributions) we show that the whole population of 8224 proteins is down-regulated in Dox-induced cells.

We visualize probability distributions again in the Figure below, but this time we analyzed both proteomic data and matching transcriptomic data.

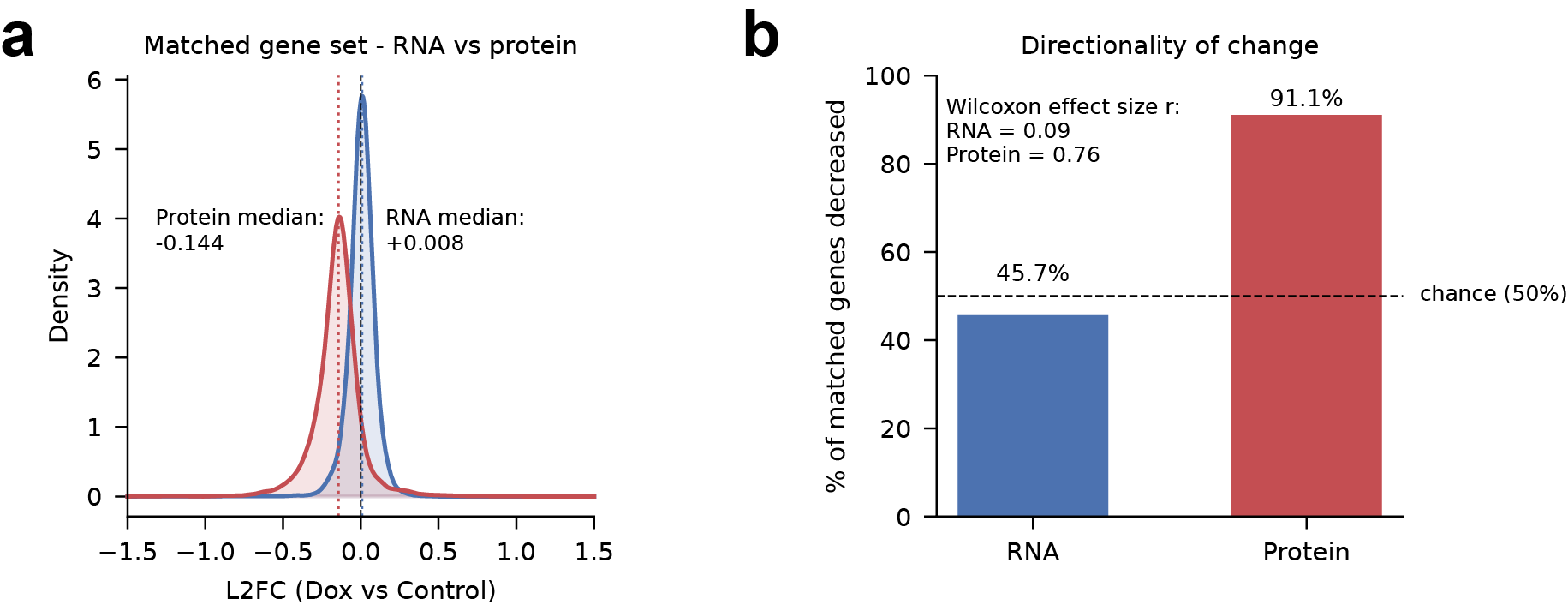

*New Fig. S6. Probability distributions of proteomes and transcriptomes (a) and directionality of changes (b) comparing Dox-induced cells (Dox) to control cells.*

At the protein level the median protein log2-fold change (L2FC) is -0.144. The mean L2FC is -0.149, which is highly similar to the median, indicating that the whole population shifts in a similar manner (panel **a**). Indeed, as shown in panel **b** the directionality of the change at the protein level is 91%. Thus, 91% of proteins is down in tubulin overexpressing cells. This effectively rules out protein degradation as a mechanism for decreased protein levels, as degradation is highly unlikely to be so aspecific.

Using our RNA-Seq dataset we analyzed the mRNA levels corresponding to the 8224 proteins in the proteomics analysis. In panel **a** of the Figure it can be seen that the median mRNA log2-fold change (L2FC) is 0.008, which is close to zero. Panel **b** shows that directionality of change is 45.7%, which is close to 50%, and is essentially what one would expect from distributions around zero. The Wilcoxon effect size (r) is 0.76 for protein, which is large, and 0.09 for mRNA, which is trivial. These data strongly suggest that transcription plays no role in the downregulation of the 8224 proteins in tubulin overexpressing cells. Thus, neither protein degradation nor transcription appear to play a role in the downregulation of proteins in tubulin overexpressing cells. We propose to add these new data to our revised manuscript and change current Fig. S6b for the new Fig. S6.

Minor comments:

Page 10." We also observed "extra-mitotic" centrioles at the onset of mitosis after Dox-treatment, indicative of mitotic arrest and aneuploidy, which eventually resolved into normal metaphase plates (Supplementary Videos 3, 4)." It is unclear whether these foci are centrioles. The SiR-tubulin reagent used in these experiment labels tubulin. It would be more accurate to describe these as "tubulin foci".

Answer:

We thank the reviewer for this comment. We will modify the text.

Page 13. "Analysis of TUBG1 reads revealed downregulation in tubulin overexpressing cells, albeit weakly (Fig. 5c)." These datapoints appear to be quite similar. Please provide a statistical analysis.

Answer:

The reviewer is correct, datapoints are quite similar. DESeq2 analysis reveals the difference in expression is minimal but significant (Dox vs Ctrl, log2fold change -0.111, p-value 0.016). These data are shown in Table S1 (sheet: DESeq2 24 hr).

**Reviewer #2**

(Note: reviewer comments are in blue, rebuttal is in black)

Summary:
This study develops an inducible HEK293F cell model that modestly overexpresses αβ-tubulin heterodimers. Using a dual expression system, the authors introduce exogenous TUBB3 and TUBA1A and combine cell biology, live-cell imaging, RNA-seq, proteomics, flow cytometry, and biochemistry to investigate the consequences of increased tubulin abundance. They suggest that, despite downregulation of endogenous tubulin transcripts, excess tubulin still accumulates in cells. This is accompanied by increased microtubule mass, changes in plus-end protein composition, defects in mitosis, and alterations in cell-cycle progression. The study further links tubulin overexpression to replication stress and broader changes in cellular physiology, including reduced abundance of mitochondrial respiratory proteins, altered translation-associated factors, and an attenuated response to hypoxic stress. In addition, the authors confirm previous observations that oxygen and nutrient deprivation reduce tubulin and microtubule abundance. Based on these findings, the authors propose that tubulin is not merely a structural component of microtubules but also contributes to the regulation of cellular homeostasis.

The possibility that tubulin abundance itself influences broader aspects of cellular physiology is conceptually interesting and warrants further investigation. At the same time, many of the mechanistic links proposed in the manuscript remain insufficiently resolved. A large number of cellular processes are associated with tubulin overexpression, but it is often unclear which phenotypes represent primary consequences of altered tubulin abundance and which are secondary responses. For instance, conclusions regarding mitochondrial dysfunction and translation attenuation are inferred from omic signatures rather than direct functional tests. Similarly, while the authors propose that altered tubulin abundance contributes to many of the observed phenotypes, the causal relationship between tubulin abundance and the downstream cellular defects is not fully established.

Despite being largely descriptive, the study has several strengths. The inducible dual-tubulin expression system is technically elegant and provides a useful platform to investigate the consequences of altered tubulin isotype composition and/or abundance. The work also indicates that relatively modest increases in tubulin levels can have measurable biological consequences, offering a possible explanation for why tubulin abundance is tightly regulated. Finally, the breadth of approaches-including transcriptomics, proteomics, imaging, and cell-cycle analyses-provides a valuable resource for the field. While many of the proposed connections remain to be mechanistically tested, the study raises a number of interesting hypotheses and highlights several promising directions for future work.

Answer:

We thank the reviewer for these positive comments.

Major comments:

Figure 1:

1. The dual expression system that the authors use to introduce exogenous tubulins is well described and characterized, representing an elegant strategy and useful tool for the field. The current version of the manuscript, however, is missing a quantitative assessment of the overall tubulin levels (endogenous + exogenous) upon doxycycline treatment (e.g., using western blot). Given that the article is centered on cellular roles of surplus soluble tubulin, the authors must quantify both the soluble and polymerized fractions of total cellular tubulin in control and doxycycline-treated samples.

Answer:

We will perform the assays requested by the reviewer and add them to a revised version of the manuscript.

2. Throughout the manuscript, the authors extrapolate the effects of TUBB3+TUBA1A overexpression to "tubulin overexpression". But how can they distinguish isotype-specific from general effects? If endogenous tubulins are massively downregulated upon the expression of exogenous TUBB3+TUBA1A, the tubulin isotype composition is presumably dramatically altered. If the authors wish to retain the claims later on in the manuscript that the observed phenotypes are associated with excess tubulin, they need to provide evidence by testing other isotype combinations. Alternatively, the authors should tone their claims to reflect the possibility that what they observe downstream of TUBB3+TUBA1A overexpression may be isotype-specific rather than general.

Answer:

Making new stable lines with other tubulin isotypes and analyzing the many downstream effects takes months and is beyond the scope of the present manuscript (see also below, *Description of analyses that authors prefer not to carry out*). Hence, we will tone down our claims by acknowledging that observations were made with one set of tubulin isotypes.

Figure 2:

1. While changes at the level of EB1, and to a lesser extent CLIP-170 upon TUBB3+TUBA1A overexpression are convincing, the claimed changes in microtubule dynamics are not supported by the data (small, but statistically insignificant trends are observed). The authors should tone down their claims and adapt the results subheadings and figure captions accordingly. Likewise, the authors state "our results convincingly show that soluble tubulin levels control +TIP composition at MT ends," which should be toned down, as the authors do not provide any experiments to probe this model (e.g., mild tubulin downregulation to restore normal protein levels and then characterize the +TIP composition).

Answer:

With respect to changes in microtubule dynamics we refer this reviewer to our answers to comments of reviewer #1 (pages 1,2). Briefly, by displaying a cumulative distribution plot in combination with appropriate tests we show that effects are statistically significant. We propose to include these data in a revised version.

We furthermore propose to perform dose-dependent doxycycline experiments (see also next comment) and test whether these cause dose-dependent +TIP composition shifts. If this is the case we maintain our current statement, if not, we will tone down this statement.

Figure 3.

1. Similar to Figure 2, the authors state that tubulin overexpression leads to mitotic defects. But this model is never challenged in rescue experiments, and the claims should therefore be toned down.

Answer:

We do not completely understand this comment. We simply turn on (or off) tubulin expression, and in our view the “rescue” experiment is the control situation, i.e. non-induced cells. To addres this criticism, we nevertheless propose to perform dose-dependent doxycycline experiments and test whether these cause dose-dependent mitotic effects.

2. The methodology that the authors chose to measure metaphase duration is inadequate, since it is difficult, if not impossible, to precisely measure metaphase duration with a tubulin label alone. A DNA label is required. Likewise, measurements of spindle length should be done in living, but not fixed cells. These parameters can be extracted from the same movies with a tubulin and a DNA stain.

Answer:

We will perform additional experiments where we add a DNA label (in addition to a tubulin label) to more accurately measure metaphase duration (and we take along spindle length).

We do not understand the claim of the reviewer that “measurements of spindle length should be done in living, but not fixed cells”. We and others have measured spindle length in fixed cells with immunostaining (PMID: 38117947, PMID: 40353778; PMID: 25568341), and it therefore appears to be an accepted method. Note that in these measurements we do include a DNA label.

3. The unaligned chromosomes in metaphase represent a striking phenotype associated with TUBB3+UBA1A overexpression. A close inspection of the mitotic spindle staining, however, doesn't seem to show a much denser microtubule network in dox-treated samples. This is surprising, and the authors should provide a quantification to support their model (e.g., tubulin fluorescence intensity normalized to an internal control), or at the very least discuss this paradox in the manuscript.

Answer:

We thank the reviewer for this observation and will analyze this in more detail.

Figure 4.

1. The authors convincingly demonstrate the alterations in cell cycle progression upon TUBB3+TUBA1A overexpression using flow cytometry. An orthogonal approach would strengthen this claim, while allowing the authors to mechanistically test the premature G1/S transition model that they propose. As is, the reduced G1 population could also be explained by increased G2/M population in these cells. Well-established time-resolved approaches exist for directly measuring G1/S transition timing, including FUCCI systems for live imaging and EdU pulse-chase methods for temporal cell cycle analysis. Without employing these validated methodologies, the inference of premature G1/S remains untested, leaving a key mechanistic link in the claim insufficiently supported.

Answer:

We thank the reviewer for this comment. We note that the flow cytometry data are supported both by RNA-Seq and proteomics results, as stated in the manuscript. Hence, orthogonal approaches were to some extent already performed. However, we do feel that additional evidence should be provided to support our premature G1/S transition model. Based on the suggestion of the reviewer we will perform an EdU pulse-chase experiment as orthogonal approach.

Figures 5-6.

1. The authors use next-generation sequencing to reveal transcriptional changes associated with TUBB3+TUBA1A overexpression. While these analyses appear comprehensive, it remains unclear how many of the alterations in gene expression are a direct consequence of TUBB3+TUBA1 overexpression and how many are a downstream consequence of the altered cell cycle profile. The authors should, at the very least, acknowledge this in the manuscript.

Answer:

This is actually acknowledged in the manuscript, on page 15: “These results are consistent with our FACS analysis and suggest that shifts in cell cycle fractions partly underlie gene expression differences”.

2. Similarly, the authors provide a careful proteomics profiling of cells overexpressing TUBB3+TUBA1 in both normoxia and hypoxia. But as with transcriptomics, it is unclear what the contribution of the altered cell cycle profile is in these analyses. This should, at the very least, be acknowledged in the manuscript.

Answer:

Again, this is actually stated in the manuscript, on page 18: “Metascape analysis revealed upregulation of terms associated with mitosis and G2/M transition, including the PLK1 pathway, in the Dox-induced cells, whereas G1/S-specific, and DNA repair terms were down (Fig. 6d, Table S1). These data are consistent with our FACS (Fig. 4a, b) and RNA-seq results (Fig. 5e)”.

3. The authors then leverage their omics data to propose a functional link between tubulin overexpression and translation and mitochondrial function. A limited number of targeted functional assays would substantially strengthen the manuscript and help distinguish between primary and secondary effects. For example, direct measurements of protein synthesis (e.g., puromycin incorporation or OPP labeling) would provide evidence for the proposed translation defects. Likewise, direct assessment of mitochondrial function (e.g., oxygen consumption, mitochondrial membrane potential, or ATP production) would strengthen claims regarding mitochondrial dysfunction. These experiments are standard in the field and could realistically be completed within several weeks to a few months, depending on local expertise and instrumentation. Moreover, they would test conclusions already central to the manuscript without opening entirely new directions.

Answer:

We thank the reviewer for these comments and helpful suggestions. We will perform experiments measuring translation and mitochondrial (dys)function (see also our rebuttal to reviewer #1 on pages 6 and 7), and include them in a revised version of the manuscript.

Figure 7.

1. The authors use proteomics-based approaches to characterize the partners of the overexpressed tubulins, revealing novel interactors. These data present an interesting but incomplete picture. For example, can the identified novel interactors bind any tubulin isotype, or are they specific to the TUBB3/TUBA1A used in this study? The enrichment of TIMM50 does not appear very strong. How reproducible are these data? The authors should provide quantifications from several independent biological replicates and perform statistical analyses.

Answer:

We thank the reviewer for these comments. As stated on page 22 of our manuscript: “Thus, AIFM1 and TIMM50 are consistently identified as TAPs using different purification strategies, tags, and cell lines”. For the revised version of the manuscript we will perform pull down experiments with other recombinant tubulins.

With respect to the relatively weak TIMM50 enrichment we refer to our answer to reviewer #1 on pages 5, 6 of this rebuttal. Briefly, we will perform additional pull down experiments (dual affinity purifications) to solidify our claims.

Figure 8.

While convincing, this figure largely confirms previous studies. In addition, it addresses a question that is different from the rest of the manuscript. The reviewer feels like this part could be taken out of the manuscript to reduce complexity and focus the scope of this work.

Answer:

Since reviewer #1 is of a similar opinion as this reviewer we are willing to remove the data presented in Fig. 8 in a revised version of the manuscript. We agree that this reduces complexity and focusses the scope of this work.

Minor comments:

1. There is a discrepancy between the manuscript and the methods that makes it unclear in which cells the transcriptomic analyses were done. The authors should clarify this.

Answer:

We are not sure what the reviewer means here, there is but one Dox-inducible cell line in which our own transcriptomic analyses was done. On page 37 we list the GEO numbers from which we retrieved expression data for Figure 8. Note that this part of the Methods section will be removed in a revised version, as we will not show Fig. 8 anymore.

2. The authors report "We also observed "extra-mitotic" centrioles...". However, they appear to refer to movies of SiR-tubulin-stained cells. This staining dows not allow visualization of centrioles, and the authors should revise their manuscript to either provide a clarification or correct this claim.

Answer:

We thank the reviewer for pointing out this mistake. We will correct this.

### 3. Description of the revisions that have already been incorporated in the transferred manuscript

NA.

### 4. Description of analyses that authors prefer not to carry out

**Reviewer #2**

2. Throughout the manuscript, the authors extrapolate the effects of TUBB3+TUBA1A overexpression to "tubulin overexpression". But how can they distinguish isotype-specific from general effects? If endogenous tubulins are massively downregulated upon the expression of exogenous TUBB3+TUBA1A, the tubulin isotype composition is presumably dramatically altered. If the authors wish to retain the claims later on in the manuscript that the observed phenotypes are associated with excess tubulin, they need to provide evidence by testing other isotype combinations. Alternatively, the authors should tone their claims to reflect the possibility that what they observe downstream of TUBB3+TUBA1A overexpression may be isotype-specific rather than general.

Answer:

We thank the reviewer for this comment. Making new stable lines with other tubulin isotypes and analyzing the many downstream effects takes months and is beyond the scope of the present manuscript. Hence, we will tone down our claims by acknowledging that observations were made with one set of tubulin isotypes.
